# Mechanistic investigation of mEos4b reveals a strategy to reduce track interruptions in sptPALM

**DOI:** 10.1101/475939

**Authors:** Elke De Zitter, Daniel Thédié, Viola Mönkemöller, Siewert Hugelier, Joël Beaudouin, Virgile Adam, Martin Byrdin, Luc Van Meervelt, Peter Dedecker, Dominique Bourgeois

## Abstract

Green-to-red photoconvertible fluorescent proteins repeatedly enter dark states, causing interrupted tracks in single-particle-tracking localization microscopy (sptPALM). We identified a long-lived dark state in photoconverted mEos4b that results from isomerization of the chromophore and efficiently absorbs cyan light. Addition of weak 488-nm light swiftly reverts this dark state to the fluorescent state. This strategy largely eliminates slow blinking and enables the recording of significantly longer tracks in sptPALM with minimum effort.

## Main text

Fluorescent proteins (FPs) and in particular green-to-red photoconvertible fluorescent proteins (PCFPs) have become indispensable tools for advanced imaging such as single-molecule localization microscopy (SMLM) or single-particle tracking photoactivated localization microscopy (sptPALM). Both techniques are however limited by blinking, a process in which the fluorophores stochastically enter reversible dark states. PCFPs display blinking on multiple timescales, arising from different underlying photochemical processes.^1^ Fluorescence intermittencies shorter than the typical exposure times used in these imaging methodologies (~tens of milliseconds), such as caused by intersystem crossing to the triplet state, reduce the apparent brightness of the label. Intermittencies longer than the exposure time, in contrast, can cause severe complications such as multiple counting of target molecules in quantitative SMLM and interruptions of single-molecule tracks in sptPALM.^2^ However, the mechanistic origin of long-lived dark states in PCFPs has remained unclear and there is presently no strategy to eliminate these.

We set out to investigate the nature of long-lived fluorescence intermittencies in photoconverted (red) mEos4b,^3^ one of the latest probes in a series of highly popular green-to-red PCFPs. Individual molecules of mEos4b were immobilized in a polyacrylamide (PAA) matrix, converted to the red emissive state using 405-nm illumination, and the fluorescence emission visualized in time using a sensitive widefield microscope. The single-molecule fluorescence traces (Fig. 1A) displayed reversible and long-lived intermittencies, which we identified as the blinking giving rise to interpretation difficulties in SMLM and sptPALM. Histograms of the intermittency duration (Supplementary Fig. 1) revealed the presence of at least two dark states, as previously reported in mEos2 or Dendra2^4,5^ (Supplementary Note 1). While the shorter-lived dark state was insensitive to the intensity of the employed 561-nm illumination (Supplementary Figure 2), the rate at which the longer-lived dark state returned to the emissive state increased with the illumination intensity, reaching a saturation regime at ~0.8 s^-1^ above a power density of ~ 1.5 kW/cm² (Fig. 1B), indicating a sensitivity to light. The shorter-lived dark state has previously been assigned to a radical state displaying a transiently distorted chromophore in mEos variants.^5^ With a typical duration of < ~100 ms, this state only causes fluorescence interruptions of at most a few frames, which can be handled by current processing software.^4^ In contrast, the longer-lived dark state may last for several seconds, and is thus much more problematic. Previous work had reported the existence of this state, including a hint of its light sensitivity^4,5^ but its nature remained elusive. To identify the underlying molecular mechanism, we first sought to link the single-molecule observations with ensemble-level measurements carried out at higher concentrations of mEos4b. Irradiation of mEos4b molecules immobilized in polyacrylamide (PAA) with 561-nm light did indeed reveal the formation of a long-lived non-fluorescent species (Supplementary Fig 3). The 561-nm laser power dependence of the longer-lived dark state displayed a similar trend as that observed with single molecule data, suggesting that the same state is observed, and that its lifetime is at least tens of minutes when kept in the dark (Supplementary Fig. 4). This also allowed us to capture the changes in absorption and emission spectra associated with slow blinking (Supplementary Fig. 3).

**Figure1.**
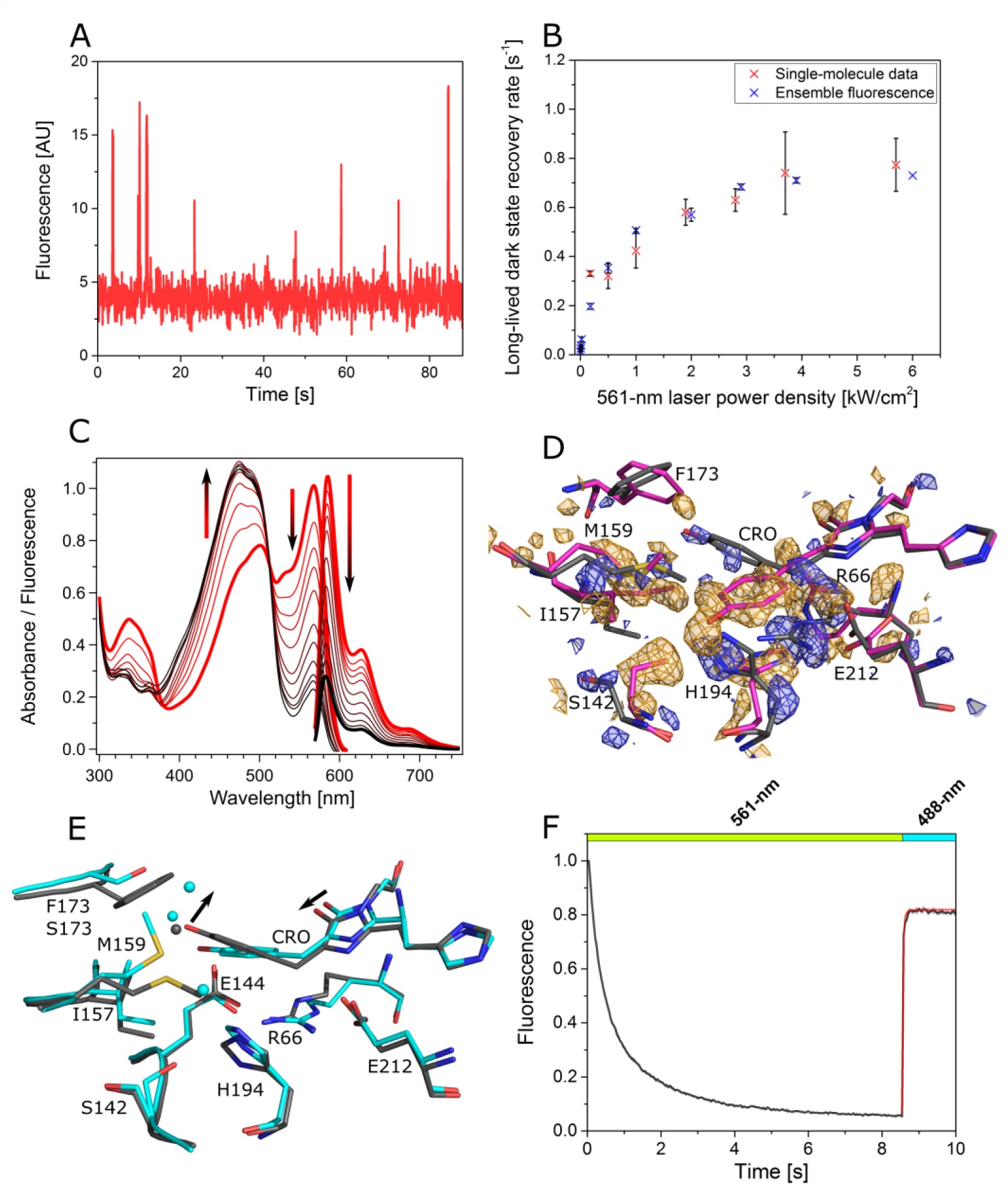
(A) Single molecule trace showing long intermittencies (seconds) under 561-nm illumination (0.5 kW/cm^2^). (B) Comparison of recovery rates from the long-lived dark state obtained from single-molecule and ensemble data on red mEos4b. Means and standard deviations from triplicate experiments are plotted. (C) Evolution (red to black, arrows) of absorption and fluorescence emission spectra from crystalline mEos4b during 561-nm illumination. (D) Comparison between the crystallographic structures of bright red mEos4b (magenta, *Red*_*on*_) and the 561-nm light-induced long-lived dark state (grey, *Red*_*off*_) in the *F_obs, illuminated_– F_obs, non-illuminated_* electron density map contoured at 3 r.m.s.d. (negative density in orange, positive density in blue). The chromophore and important residues of the chromophore pocket are shown (E) Comparison between the crystallographic structures of the *Red*_*off*_ state in mEos4b (grey) and the off-switched structure of green IrisFP (PDB ID: 2VVI). Arrows indicate the shift in imidazolinone ring and consecutive distortion of the chromophore in *Red*_*off*_ compared to IrisFP. (F) Ensemble photoswitching of red mEos4b under 561- and 488-nm light (80 and 5 W/cm^2^, respectively). Upon illumination at 488 nm, fluorescence recovery occurs in a few hundreds of milliseconds. This recovery can be fitted with a monoexponential model (red line), yielding a rate of 45 s^-1^.

Next, we set out to visualize the underlying structural change using kinetic crystallography.^6^ To this aim we generated crystals of red-state mEos4b and illuminated them at 561 nm. The resulting changes in spectra (Fig. 1C) were very similar to those observed in PAA, supporting that the longer-lived dark state could be accumulated *in crystallo* (Supplementary Note 2). Hence, we selected a single crystal and illuminated one part at 561 nm, leaving the other part unexposed. X-ray diffraction on the flash-frozen crystal revealed that both parts contained a mixture of the mEos4b green and red forms (Supplementary Fig. 5), but that the illuminated part additionally contained a fraction of mEos4b in the longer-lived dark state (Fig. 1D). Using difference refinement, we selectively extracted the structure of this species, denoted *Red*_*off*_ (Online Methods; Fig. 1D and Supplementary Fig. 6). The chromophore in *Red*_*off*_ adopts a strongly bent or “frustrated” *trans* configuration, being forced away from a more canonical *trans* configuration (Fig. 1E) to avoid steric clashes with the nearby Ile157 (Supplementary Fig. 7), and is also found to be highly dynamic (Supplementary Note 3, Supplementary Fig. 8). The structural signature of the *Red*_*off*_ dark state is reminiscent of the photochromism observed in reversibly switchable fluorescent proteins.^7,8^ In these FPs, illumination in the main absorption band populates a nonfluorescent, spectroscopically blue-shifted state, corresponding to a protonated *trans* chromophore. The fluorescent state can then be efficiently recovered by illumination in this newly-arisen band. In mEos4b the dark state absorbs maximally at ~475 nm and we indeed observed prompt recovery of red fluorescence (~80%) under mild 488-nm illumination (Fig. 1F). This finding suggested a straightforward and readily-accessible way to suppress long-lived intermittencies in SMLM and sptPALM experiments, by weakly illuminating in the absorption band of the *Red*_*off*_ chromophore either continuously or in a pulsed manner to avoid residual background from excited green molecules (Fig. 2A). Indeed, weak illumination at 488-nm of mEos4b molecules embedded in PAA revealed a clear suppression of the longer intermittencies by about one order of magnitude (Fig. 2B, Fig. 2C), leading to an over tenfold increase in the number of molecules that displayed on-times larger than 3 s. Similar results were obtained in fixed HeLa cells expressing mEos4b in the cytosol (Supplementary Fig. 9), as well as *in vitro* using other mEos variants (Supplementary Fig. 10). At the used power densities, the addition of 488-nm light did not change the total number of photons that could be detected from a single red mEos4b molecule before irreversible photodestruction (Supplementary Fig. 11), nor did it cause significant photobleaching of green mEos4b molecules (Supplementary Note 4, Supplementary Fig. 12). Moreover, we observed a slightly higher rate of green-to-red mEos4b photoconversion that reduced the need for additional 405-nm light (Supplementary Fig. 13).^9^ Recent efforts have focused on increasing single-particle track lengths either using organic dyes in reducing/oxidizing buffers,^10^ multiple copies of FPs in combination with sparse labelling^11^ or FRET-enhanced photostability.^12^ However none of these methods offer the generality of using single FPs in live-cell compatible environments. We reasoned that our strategy could provide longer and more informative single-molecule tracks in standard sptPALM experiments. To this aim, we explored the diffusion behaviour of Microtubule-Associated Protein 4 (MAP4) fused to mEos4b and expressed in Cos7 cells.^13^ Visual inspection of raw sptPALM data suggested that much longer mEos4-MAP4 tracks could be obtained when a small amount of 488-nm light (4.8 W/cm²) was added (Fig. 2D, Supplementary Fig. 14, Supplementary Movie 1). The quantitative evaluation of the data indeed indicated that addition of cyan light resulted in a significant increase of the average track length by 54 % and in the appearance of over 15-fold more tracks lasting longer than 2 s (Fig. 2E). These long tracks revealed a heterogeneity in MAP4 dynamics, likely due to switching of individual molecules between at least three diffusion states (unbound, bound immobile and bound mobile along the microtubules) (Fig. 2F). The many more tracks displaying a large number of diffusion state changes in the presence of weak 488-nm light (Fig. 2G) opens the door to statistically significant investigations of the complex MAP4 dynamics. Importantly, the addition of 488-nm light did not have a significant influence on the observed diffusion coefficients (Supplementary Fig. 15).

**Figure 2.**
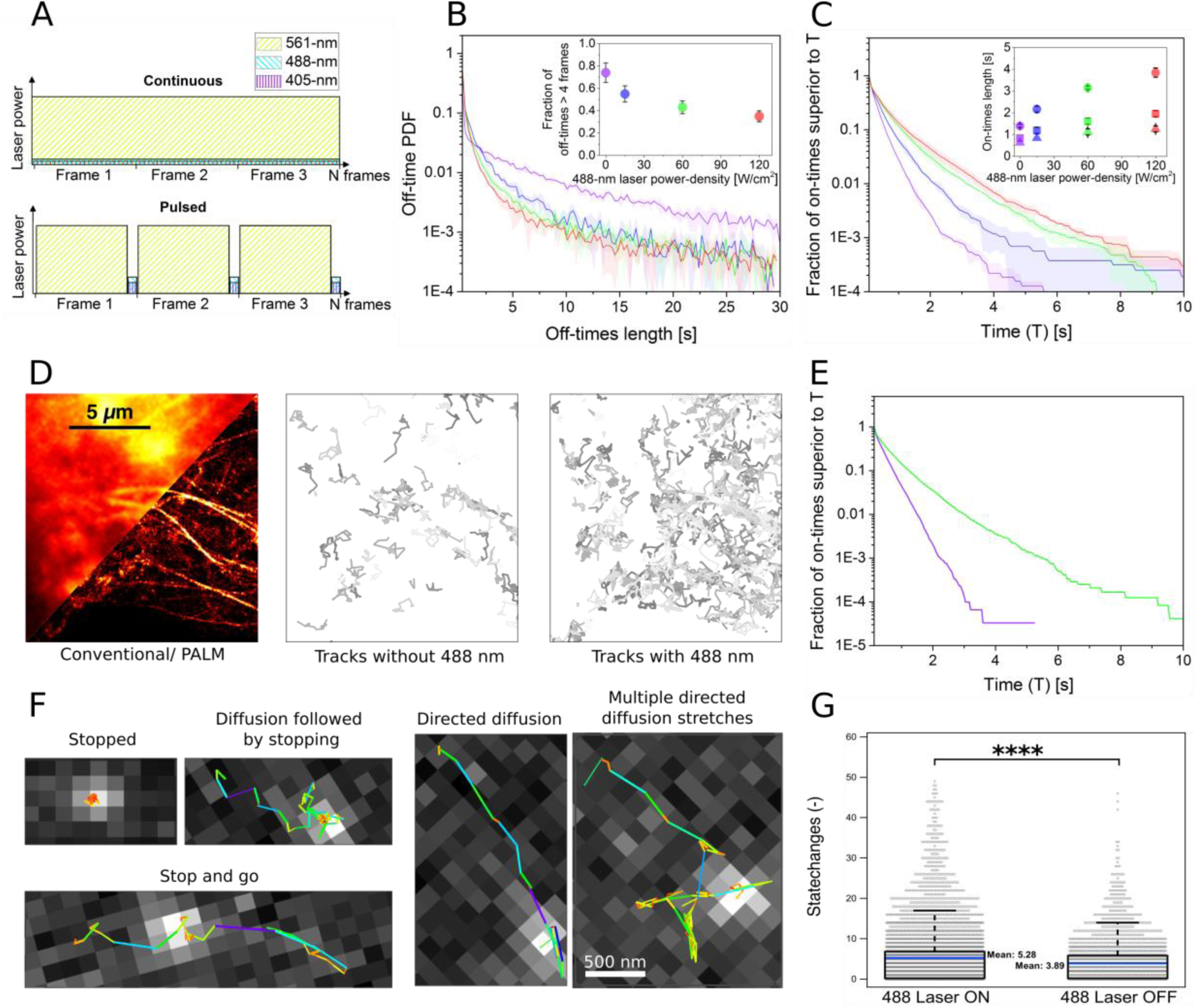
(A) Proposed illumination schemes for suppression of long-lived intermittencies in sptPALM. Weak 488-nm light is added to the standard readout 561-nm light, either continuously for simplicity or in an interleaved manner to minimize background fluorescence during frame exposure. (B) Intermittency histograms for red mEos4b under sole 561-nm light (0.5 kW/cm^2^, violet curve), or combined 561- and 488-nm light (15, 60 or 120 W/cm^2^ during 1/10^th^ of the frametime, blue green and red curves, respectively). PDF: Probability Density Function. The fraction of intermittencies longer than 4 frames decreases with additional 488-nm illumination (Inset). (C) Cumulative distribution of single-molecule on-times longer than a given time (corrected for intermittencies shorter than 4 frames) under sole 561-nm light or combined 561- and 488-nm light. Long on-times become more frequent upon addition of 488-nm light. The inset shows the minimum on-times displayed by a given fraction of the data (1%, circle; 5%, square; 10%, triangle), as a function of the applied 488-nm laser power density (D) Imaging of COS-7 live cells labeled with MAP4-mEos4b. Left: conventional image and PALM reconstruction, middle and right: single particle trajectories longer than 1 s in the absence and presence of 488-nm light (4.8 W/cm²). (E) Cumulative distribution of track lengths from all MAP4-mEos4b tracking experiments, in the absence (violet) or presence (green) of additional 488-nm illumination. (F) Panel of representative diffusion patterns identified amongst the large set of long tracks that become visible upon addition of 488-nm light, color coded according to step length, overlaid on raw data from randomly chosen single frames. (G) Swarm and box plots of the amount of state changes extracted by the HMM-Bayes software in the presence (left) and absence (right) of 488-nm light (*: χ^2^ test with p << 0.01).

Although addition of 488-nm light eliminated a large fraction of the long intermittencies in mEos4b, the suppression was not complete and resulted in an increased number of short intermittencies (Fig 2C). These observations can be explained by the tentative photophysical scenario depicted on Supplementary Fig. 16 and Supplementary Note 5, involving an additional non-emissive state, which would also account for the saturation behavior observed in Fig 1B. Importantly, although *Red*_*off*_ can be excited by 405-nm light relatively efficiently (Supplementary Fig. 17), illumination at this wavelength is constrained by the desirable rate of green-to-red photoconversion and generates phototoxicity.

In conclusion, our work has revealed that long-lived intermittencies observed in mEosFP-like fluorophores predominantly result from photochromism as seen in reversibly switchable or biphotochromic FPs^14^ and can be drastically reduced by simple structure-guided “rational microscope engineering”. Since a 488-nm light source is present on most instruments capable of sptPALM, weak illumination at this wavelength is likely to be readily applicable, providing a strong improvement in data quality with a minimum of effort. Our blinking suppression strategy should be similarly useful in increasing the accuracy of molecular counting in quantitative SMLM imaging.

## Supporting information

## Authors contributions

DB initiated research. EZ, PD and DB designed the project. EZ, VA, MB and DB carried out mechanistic investigations. DT performed in vitro single-molecule and ensemble experiments with contributions from EZ and DB. VM, SH and PD performed sptPALM experiments. DT and JB performed fixed cells experiments. LvM assisted in the crystallographic analysis. EZ, DT, PD and DB wrote the manuscript with input from all authors.

## Acknowledgement

This work was supported by the Agence Nationale de la Recherche (ANR-17-CE11-0047-01) and used the M4D imaging platform of the Grenoble Instruct-ERIC Center (ISBG: UMS 3518 CNRS-CEA-UGA-EMBL) with support from FRISBI (ANR-10-INBS-05-02) and GRAL (ANR-10-LABX-49-01) within the Grenoble Partnership for Structural Biology (PSB). E.D.Z. thanks the Research Foundation Flanders (FWO) for a doctoral fellowship and travel bursary. S. H. thanks the Research Foundation Flanders (FWO) for a postdoctoral fellowship. We also acknowledge support from the European Research Council for ERC Starting Grant NanoCellActivity and the Research Foundation Flanders (FWO) for grant G0B8817N (both to P.D.). The authors thank the staff of beamline ID-23-1 from the European Synchrotron Radiation Facility (ESRF, Grenoble, France).

## Methods

### Cloning, protein expression and purification

mEos4b was expressed from a pRSETb plasmid (Invitrogen) in JM109(DE3) cells (Promega, Leiden, The Netherlands) in 1l LB containing ampicillin. After four days of growth at 21ºC, the cells were harvested by centrifugation (20 minutes at 5000 rpm), resuspended in TN buffer (100 mM Tris, 300mM NaCl, pH 7.4) and lysed using a French pressure cell press (1150 psi). After spinning down the cell debris (20 minutes at 8500 rpm), a Ni-NTA affinity chromatography was carried out using a HisTrap FF crude column (GE Healthcare, Diegem, Belgium) coupled to an Akta Prime system (GE Healthcare, Diegem, Belgium) and TN buffer supplemented with 500 mM imidazole. The sample was then loaded onto a HiLoad Superdex 200pg 16/600 column (GE Healthcare, Diegem, Belgium) coupled to an Akta Purifier 10 system (GE Healthcare, Diegem, Belgium) for size-exclusion chromatography. Elution was performed using 0.1xTN buffer. Finally, the protein was concentrated using a Vivaspin 10K MWCO column.

CD86-mEos4b was cloned by replacing mEos2 by the gene coding for mEos4b in the CD86-mEos2 construct (Addgene # 98284),^1^ using the Gibson Assembly^®^ Master Mix (NEB, Ipswich, Massachusetts, United States). The MAP4-mEos4b construct was generated by replacing mCherry for the mEos4b gene in the mCherry-MAP4-N-10 construct, purchased at Addgene (plasmid # 55076), using the AgeI and NotI restriction sites.

### Mutagenesis and screening

Site saturation mutagenesis was performed using a modified QuikChange protocol^2^. The mEos4b point mutants were expressed in JM109 cells. Screening for altered blinking/ photoswitching behavior was performed on a home-built colony imaging system containing a 300 W Xenon lamp (MAX-302, Asahi Spectra)^3^. Green colonies were exposed for 60 min to light passing through a 480/40 bandpass filter and then for 10 min to light passing a 400/30 bandpass filter, and this illumination cycle was repeated twice. Every minute an image was acquired through a 530/40 bandpass filter on an EMCCD camera (Cascade 512B, Photometrics).

Colonies showing altered blinking/photoswitching behavior were selected and further investigated using an inverted microscope (Olympus IX71) containing a Sola Light Engine (Lumencor)^3^. 200 μl cell cultures expressing the mutant proteins were subjected twice to 5 s cyan light (50 and 75 %) and 5 s violet light (5%). Then the proteins were photoconverted using 30 s violet light (100%) and subsequently exposed twice to yellow-green (5 s, 50 and 75 %) and cyan light (5 s, 5%). Every second an image was captured on an EMCCD camera (iXon, Andor).

### Crystallization

Green mEos4b crystals were grown in sitting drops in which 1 µl of protein was mixed with 1 µl precipitant (100 mM HEPES pH 7.5 and 30 % PEG 1000) equilibrated against 100 µl of precipitant. Rod-shaped crystals (0.1 × 0.1 × 0.5 μm^3^) grew in a few weeks. Subsequently, crystals were exposed to sunlight to photoconvert mEos4b to the red state.

### Cell culture and transfection

#### mEos4b fused to CD86 in HeLa cells

HeLa cells were grown in DMEM growth media containing 10% FBS and maintained at 37°C in 5% CO2. Cells were seeded in a tissue-culture treated 6 well plate and transfected using X-tremeGENE 9 (Sigma-Aldrich, St. Louis, Missouri, United States). Three days later, cells were trypsinated and replated in a sticky-Slide VI 0.4 (Ibidi GmbH, Martinsried, Germany) mounted with a 24X50mm #1 Menzel coverslip (Fisher scientific, Illkirch, France) coated with fibronectin (Sigma-Aldrich, St. Louis, Missouri, United States). 5 hours later, cells were rinsed with PBS and fixed with a mix of 4% formaldehyde (stock 16%, Thermofisher, Waltham, Massachusetts, United States) and 0.1% glutaraldehyde (stock 25%, Sigma-Aldrich, St. Louis, Missouri, United States) in PBS. Cells were then washed and kept in PBS for imaging.

#### mEos4b fused to MAP4 in COS-7 cells

COS-7 cells were grown in DMEM growth media containing 10% FBS and maintained at 37°C in 5% CO2. Cells were seeded in sterile 35 mm glass-bottom imaging dishes and transfected using FuGENE (Promega) following the manufacturer’s protocol. After incubation for 24 h, the cells were washed twice with Hank’s Balanced Salt Solution (HBSS, 14065, Life Technologies) buffer for live cell imaging and subsequently imaged in HBSS buffer at room temperature. For fixation the cells were washed twice with PBS and subsequently fixed with 4% Paraformaldehyde/ 0.2%Glutaraldehyde for 10 min at 37°C in 5% CO2.

### Microscopy experiments

Ensemble and single-molecule *in vitro* and CD86 data were acquired on a home-built PALM setup based on an Olympus IX81 inverted microscope equipped with a 100× 1.49 NA oil-immersion apochromatic objective lens (Olympus, Japan). Widefield illumination was achieved by focusing the diode-pumped solid state 405-nm (CrystaLaser, USA), 488-nm (Spectra Physics, USA) and 561-nm (Cobolt Jena, Sweden) laser beams to the back focal plane of the objective. Intensities of laser illuminations at the sample were tuned by an acousto-optical tunable filter (AOTF, AA Opto Electronic, France). Fluorescence images were acquired with an Evolve 512 back-illuminated EMCCD camera (Photometrics, USA) controlled by the Metamorph software (Molecular Devices, USA).

#### mEos4b immobilization for in-vitro experiments

Purified mEos4b was diluted to either micromolar (ensemble) or nanomolar (single-molecule) concentrations in a 15% polyacrylamide (PAA) gel. The mEos4b-containing PAA was then spread to form a uniform thin layer on a coverslip thoroughly cleaned using a UV Ozone Cleaning System (HELIOS-500, UVOTECH Systems), and let to harden at room temperature for 5 minutes.

#### In-vitro single-molecule experiments

PALM data were collected using an exposure time of 70 ms, under continuous 561-nm illumination only at different laser power-densities (0.2, 0.5, 1, 2, 3, 4 and 6 kW/cm^2^) or under 561-nm illumination at 0.5 kW/cm^2^ interleaved with pulsed 488-nm illumination for 10 ms per frame at different power-densities (0, 15, 60 and 120 W/cm^2^). Illumination at 488-nm was applied between camera exposures to avoid any contribution of the green mEos4b molecules to the recorded signal. Experiments were performed in triplicate.

#### Determination of on-switching rates from single-molecule experiments

Data were processed as described in ref ^4^ Briefly, after localization of the single molecules using the Thunderstorm plugin for ImageJ,^5^ their fluorescence traces were computed and off-times were extracted and plotted as a histogram. Although off-times histograms could be precisely fit with a three-exponential model (Supplementary Fig. 1B, Supplementary Note 1), the extracted rates could not be directly related to the rates defining the kinetic scheme of Supplementary Fig. 16. Therefore, we used the single-molecule fluorescence traces to reconstitute fluorescence decay curves, by counting the number of fluorescent and non-fluorescent molecules at each time-point after activation. The curves obtained were similar to those obtained from ensemble-level experiments, and were fitted using the same kinetic model (see below). The first point of the decay curves was however not taken into account, as we found that this point was affected by residual false positive localizations detected on single frames.

#### In-vitro ensemble fluorescence

For ensemble measurements, samples were first illuminated with 405-nm light (1 W/cm²) for 1 min to prompt photoconversion. To assess the photoswitching behavior of red mEos4b (Supplementary Fig. 17), alternating 561-nm light (25 W/cm²) and 405-nm light (1 W/cm^2^, 60 seconds off, 30 seconds on) were used. To test the fluorescence recovery induced by illumination at 488-nm (Fig. 1F), red mEos4b was alternately illuminated with 561-nm light (80 W/cm^2^, 8.5 seconds) and 488-nm light (5 W/cm^2^, 1.5 seconds). To measure the dependence of the on-switching rate on the 561-nm laser power (Fig. 1B), single off-on switching cycles were acquired under the same conditions as described above for single-molecule experiments, except that measurements at much lower power-densities of the 561 nm laser could be performed, as single-molecule sensitivity was not required. Experiments were performed in triplicate.

#### Determination of on-switching rates from in-vitro ensemble experiments

Analysis of the off-on switching cycles was carried out on a small, homogenous region of the sample. For each applied 561-nm laser power, the corresponding fluorescence decay profile was fitted with a kinetic model including photobleaching and two reversible dark-states. The (thermally driven) recovery rate of the short-lived dark state was fixed to 16 s^-1^ as fitted on the single-molecule data, to prevent fitting of several non-independent constants. This value was also found consistent with ref ^6,7^. The fitted recovery rate then corresponds to an effective, slow on-switching rate whose value is affected by the power-dependent build-up of a potential extra dark state *Dark_2_* (Supplementary Fig. 16)

#### In-crystallo ensemble fluorescence

To assess the photoswitching behavior of crystalline mEos4b, crystals were mounted between two coverslips in a drop of mother liquor and placed on the PALM microscope. The same illumination scheme was used as described above for the *in-vitro* ensemble fluorescence experiments, using 70 W/cm² of 561-nm laser light for off-switching and 0.3 W/cm² of 405-nm light for on-switching. The higher power density of 561-nm light and lower power density of 405-nm light as compared to that used for ensemble experiments in PAA was chosen to compensate for the high optical density of crystals at 561 nm (Supplementary note 2). Data analysis was carried out on a small region of the crystal. Experiments were performed in triplicate.

#### sptPALM measurements and analysis

##### SPT acquisition

Imaging of the transfected cells was done on a commercial Cell TIRF DX83 microscope (Olympus) using a 100x (NA 1.49) oil-immersion objective and EMCCD camera (Hamamatsu, C9100-23B EM X). The power of the 405-nm laser was set at 0.4 – 0.6 % (0.3 – 0.6 mW/cm^2^) in the absence of 488-nm light and 0 - 0.5% (0 - 0.34 mW/cm^2^) in its presence, that of the 488-nm laser at 35% (4.8 W/cm^2^, cw mode) and that of the 561-nm laser at 35% (43 W/cm^2^) in a HILO (Highly Inclined and Laminated Optical) illumination configuration. The dose used for the 488-nm laser matches the one used during in-vitro experiments, where pulsed light was applied. Because the 488-nm light also resulted in residual green-to-red conversion (Supplementary Fig. 13), the intensity of the 405 nm light was reduced to maintain the approximate same number of active molecules per frame. The exposure time was 40 ms with an EM gain of 500. On each cell we recorded two datasets, one with and one without 488-nm light. On half of the cells, the dataset with 488-nm light was recorded first, while on the other half the dataset without 488-nm light was recorded first. The appearance of longer tracks was independent of the order in which these datasets were acquired.

##### SPT analysis and statistical relevance

For track visualization we employed the Localizer software package,^8^ implemented in Igor Pro (WaveMetrics). For analysis of the localizations, the default settings were used with a generalized likelihood ratio test (GLRT) insensitivity of 25 for segmentation and a standard deviation of the PSF of 1.6 pixels using the Gaussian PSF model (160 nm pixel size). For analysis of the track lengths the parameters jump distance was set to 4 pixel, blinking frames to 4, and the minimum track length to 3.

For quantitative analysis of the diffusion coefficients we applied the HMM Bayes software package^9^, which builds on output from the UTrack software package.^10^ UTrack was applied with the same settings as for the Localizer-based analysis. HMM Bayes was employed with default settings to estimate diffusion coefficients corresponding for each observed track. Only tracks longer than 25 frames were used in the HMM-Bayes evaluation to speed up the analysis. All calculations, including all statistical tests (χ^2^ tests), were performed in Matlab R2018a (MathWorks).

##### Spectroscopic characterization and Crystal preparation

Ensemble and in crystallo absorbance and fluorescence spectra were acquired on a custom-build microspectrophotometer (CAL(AI)^2^DOSCOPE)^11^ with the sample placed at the focus of two opposite 15× mirror objectives (Edmund optics, France). A 561-nm laser (Crystal laser, USA), 405-nm laser (New Industries Optoelectronics Tech, PRC) and UV-VIS light source were coupled into the optical path using optical fibers. Two AvaSpec ULS2048 spectrophotometers (Avantes, The Netherlands) collected the transmitted or emitted light while the sample surface could be visualized on a color detector (uEye GigE UI-52405E, IDS, Germany). Every 2 s an absorbance and fluorescence spectrum was acquired using respectively the white light source and the actinic light as excitation light source. The illumination and detection scheme were controlled by a 9518PG pulse generator (Quantum composers, USA) and the Avasoft software (Avantes, The Netherlands). Data analysis was performed in Igor Pro (WaveMetrics).

##### In vitro ensemble experiments

*In vitro* ensemble measurements were performed at room temperature on cover slips bearing mEos4b embedded in 1 % PVA in PBS. Photoconverted samples were illuminated for 22.5 s with 561-nm light (28 W/cm^2^) and 7.5 s with 405-nm light (0.2 W/cm^2^).

##### In crystallo experiments

Needle-shaped crystals were mounted between two coverslips in a hanging drop of mother liquor. All experiments were performed at room temperature. One part of a photoconverted red crystal was illuminated with 561-nm light (34 W/cm^2^) during 43.5 s, leaving the other part unexposed. For the control experiment, a photoconverted red crystal was illuminated in a similar manner and subsequently during 18 s with 405-nm light (1.0 W/cm^2^). Incomplete photoswitching was deliberately chosen to avoid the built-up of irreversibly photobleached molecules within the crystal.

Of note, we used 405-nm rather than 488-nm light for back switching of *Red*_*off*_ using our microspectrophotometer. Both wavelengths in fact induce back switching. Whereas the use of significant 405-nm light to promote back switching would be problematic in single molecule experiments due to coupling with photoconversion, this is not the case in ensemble experiments were samples are photoconverted beforehand.

As the thermal recovery rate from the *Red*_*off*_ dark state is slow, sufficient time was available to safely trap the proteins in this dark state by soaking them immediately after illumination for a few seconds through a cryoprotectant solution (10% glycerol in 100 mM HEPES pH 7.5 and 30 % PEG 1000), before flash freezing and storing in liquid nitrogen. Of note, the time required for this procedure (< 1 min) likely allowed molecules trapped in the *Dark_2_* state to thermally relax to *Red*_*on*_.

#### X-ray diffraction data collection and analysis and structure refinement

X-ray diffraction data of green mEos4b and the illuminated and non-illuminated parts of the red mEos4b crystals were collected at the ID-23-1 beamline of the European Synchrotron Radiation Facility (ESRF, Grenoble, France), under a nitrogen stream of 100 K with an X-ray wavelength of 1 Å. The data was indexed, integrated and scaled using XDS and XSCALE v. October 15, 2015.^12^

The structure of mEos4b in its green state was solved via molecular replacement using Phaser^13^ using chain A of mEosFP (PDB ID 1ZUX^14^) as a model. Structure refinement was carried out using phenix.refine v.1.11^15^ and COOT v. 0.8^16^ and consisted of bulk-solvent and anisotropic scaling, individual coordinate and isotropic B-factor refinement and occupancy refinement for residues in alternative conformations. After a first round of refinement, the chromophore and the mutations compared to the phasing model were clear in the *F*_*obs*_-*F*_*calc*_ difference map. Water molecules were added to the model if they appeared in the *mF*_*o*_-*DF*_*c*_ difference map contoured at 3.0 r.m.s.d., had acceptable B-factors and were within reasonable distance from chemically relevant groups. The crystallographic dictionary file for the chromophore was generated using eLBOW^17^ and manually adapted.

The structure of mEos4b in its red state (being the non-illuminated part of the red crystal) was determined starting from the green state model. After one cycle of rigid body refinement, the protein backbone break characteristic of the green-to-red photoconversion was clear in the *F*_*obs*_-*F*_*calc*_ electron density map, albeit incomplete, indicating a mixture of the green and red form of mEos4b. Restrained structure refinement was performed as mentioned for the green state above. The red and green state chromophore occupancies converged towards a 60:40 red:green ratio, which is in accordance to the crystal’s absorption spectrum. Again, crystallographic library files for the chromophore and non-standard amino acids were generated using eLBOW and manually changed. With the final model, an ensemble refinement^18^ was carried out using the default parameters except for the fraction of atoms included in the TLS fit being 0.9.

In the 561-nm illuminated part of the red crystal, a large fraction of mEos4b molecules were still found in the bright red or non-photoconverted green state, as expected. To be able to highlight features specific to the *Red*_*off*_ dark state, we calculated a Bayesian q-weighted19 *F*_*obs,illuminated*_– *F*_*obs,non-illuminated*_ electron density difference map, using the red state model to calculate phases. Based on an occupancy of *Red*_*off*_ established at 25%, extrapolated structure factor amplitudes corresponding to the pure *Red*_*off*_ dark state were then calculated as described in Coquelle et.al,^20^ and these were used in further structure refinement of *Red*_*off*_, for which we applied the same strategy as described above for the *Red*_*on*_ red state, albeit starting with a rigid body refinement of the red state model.

To ensure that the dark-state structure obtained mostly originates from *Red*_*off*_ and not from irreversibly photobleached states, we repeated the same crystal illumination procedure except that before flash-cooling we applied additional 405-nm light to induce recovery of the bright state. Diffraction data of both the illuminated and non-illuminated sides of the crystal were acquired and a q-weighted *F*_*obs, illuminated 561nm+405nm*_– *F*_*obs, non-illuminated*_ electron density difference map was calculated. This map showed no significant difference between both sides (Supplementary Fig. 19C).

All data collection and refinement statistics can be found in Supplementary Table 1 and Supplementary Table 2.

