## Supplementary material for "Mechanistic investigation of mEos4b reveals a strategy to reduce track interruptions in sptPALM"

| Section | Page |
|---|---|
| Supplementary Figures | |
| Supplementary Figure 1: Fitting of mEos4b intermittency histograms. | 3 |
| Supplementary Figure 2: Recovery rate from the shorter-lived dark-state. | 4 |
| Supplementary Figure 3: Evolution of absorption and fluorescence emission spectra from PVA-embedded mEos4b during 561-nm illumination. | 5 |
| Supplementary Figure 4: Kinetic of red mEos4b thermal recovery after off-switching. | 6 |
| Supplementary Figure 5: Green and photoconverted red state crystal structure of mEos4b. | 7 |
| Supplementary Figure 6: Electron density maps of the bright and long-lived dark state of red mEos4b. | 8 |
| Supplementary Figure 7: The dark state chromophore adopts a frustrated *trans* conformation. | 9 |
| Supplementary Figure 8: Ensemble refinement of the bright and dark state mEos4b chromophore. | 9 |
| Supplementary Figure 9: Intermittency histogram of mEos4b fused to the CD86 receptor in a fixed HeLa cell. | 10 |
| Supplementary Figure 10: Increase of on-times in the presence of 488-nm light for different popular FPs used in PALM. | 11 |
| Supplementary Figure 11: Photon budget of single red mEos4b molecules under combined 561- and 488-nm illumination. | 12 |
| Supplementary Figure 12: Illumination at 488-nm during a PALM experiment induces minor photobleaching of green mEos4b. | 13 |
| Supplementary Figure 13: Additional photoconversion due to 488-nm illumination. | 14 |
| Supplementary Figure 14: Influence of 488nm illumination during single particle tracking. | 15 |

| | |
|---|---|
| Supplementary Figure 15: Probability density function of the log10-transformed diffusion coefficients from MAP4-mEos4b tracking experiments. | 16 |
| Supplementary Figure 16: Tentative photophysical scheme for red mEos4b. | 17 |
| Supplementary Figure 17: PAA embedded and *in crystallo* red mEos4b reversible photoswitching. | 18 |
| Supplementary Figure 18: Superposition of the dark chromophore and its environment in red mEos4b and reversibly switchable red fluorescent proteins. | 19 |
| Supplementary Figure 19: Reversibility of the dark state trapped in crystalline mEos4b. | 20 |
| Supplementary Figure 20: Simulation of the rate saturation effect upon 561-nm laser power titration. | 21 |
| Supplementary Tables | |
| Supplementary Table 1: Data collection and refinement statistics of the bright and dark-state crystal structures of mEos4b. | 22 |
| Supplementary Table 2: Data collection and refinement statistics of control crystal structures of mEos4b. | 23 |
| Supplementary Table 3: Average chromophore torsion and bond angles. | 24 |
| Supplementary Table 4: Rate constants used for simulation of ensemble level experiments. | 24 |
| Supplementary Notes | |
| Supplementary Note 1: Fitting of fluorescence intermittencies histograms | 25 |
| Supplementary Note 2: Ensemble-level dark-state accumulation in solution and *in-crystallo*. | 25 |
| Supplementary Note 3: Structural basis for the frustrated trans conformation of the mEos4b chromophore in the long-lived dark state. | 25 |
| Supplementary Note 4: Side effects of 488-nm illumination. | 27 |
| Supplementary Note 5: Photophysical scheme of red mEos4b, and hypothesis of an additional dark state Dark2. | 27 |

**Supplementary Figure 1.**

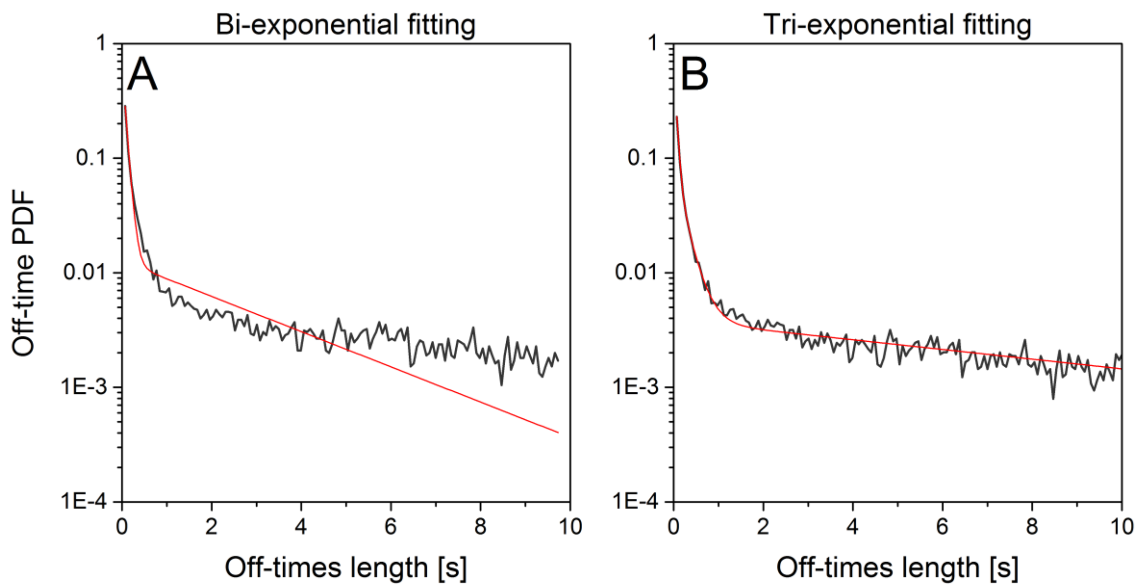


**Fitting of mEos4b intermittency histograms.**

(A) Fitting of mEos4b intermittency histogram (black line) with a bi-exponential model (red line). The model fails to accurately fit the long intermittencies (> 1 second). (B) Fitting of the same histogram with a tri-exponential model (red line). This model allows accurate fitting of the whole intermittency histogram. However, while the rapid phase can be safely associated to the recovery rate of the shorter-lived dark state, the two slower phases cannot be simply related to the recovery rate of the longer-lived dark state, justifying the approach taken to estimate this rate (see Online Methods).

**Supplementary Figure 2.**

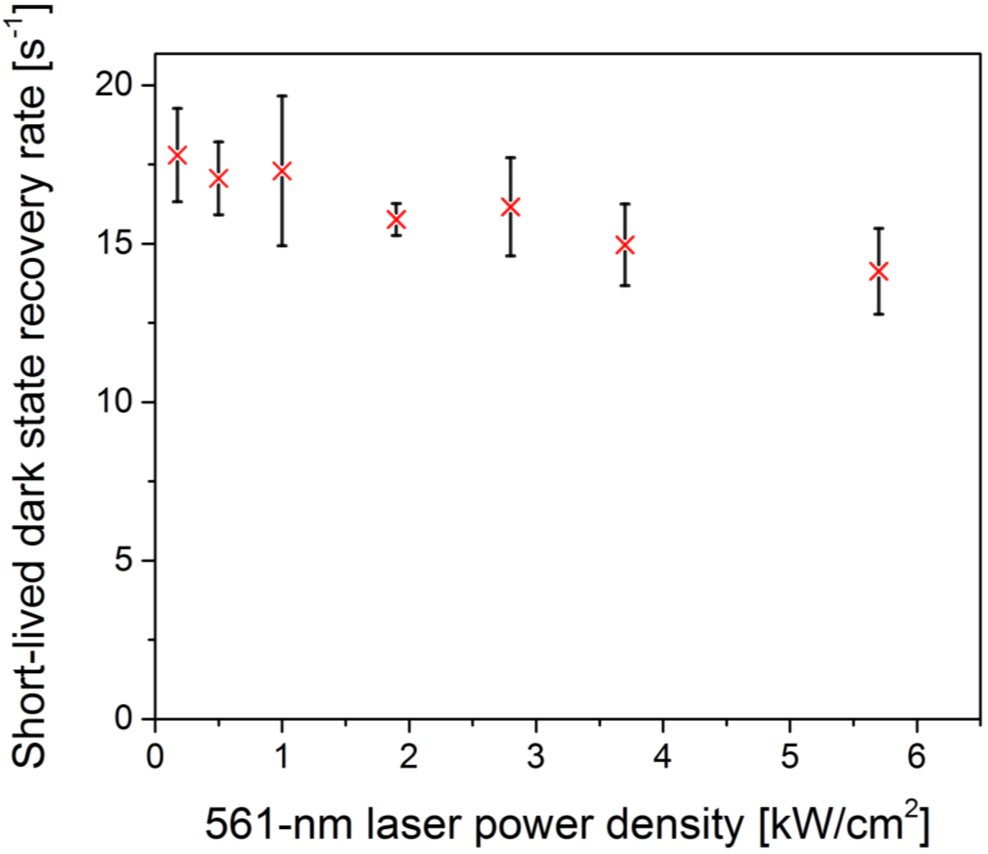


**Recovery rate from the shorter-lived dark-state.**

Recovery rates from the shorter-lived dark-state could be retrieved at different 561-nm laser power densities from the tri-exponential fit of the intermittency histograms (Supplementary Fig. 1B). The rates do not significantly vary with the increase in laser power-density, indicating that the recovery from this dark-state is essentially 561-nm light-independent. Error bars correspond to standard deviations from triplicate experiments.

**Supplementary Figure 3.**

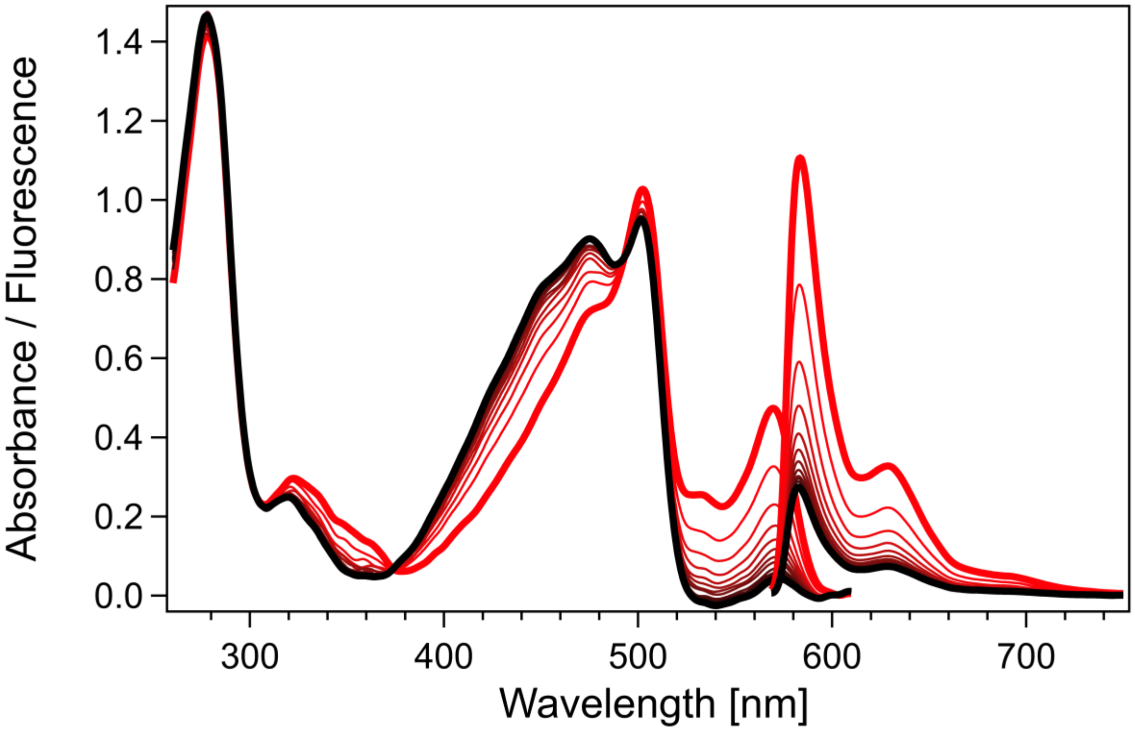


**Evolution (red to black) of absorption and fluorescence emission spectra from PVA-embedded mEos4b during 561-nm illumination.**

Upon illumination at 561 nm, the absorbance peak of the anionic red chromophore (570 nm) decreases, while a new absorbance peak appears at ~475 nm, corresponding to a long-lived non-fluorescent state.

**Supplementary Figure 4.**

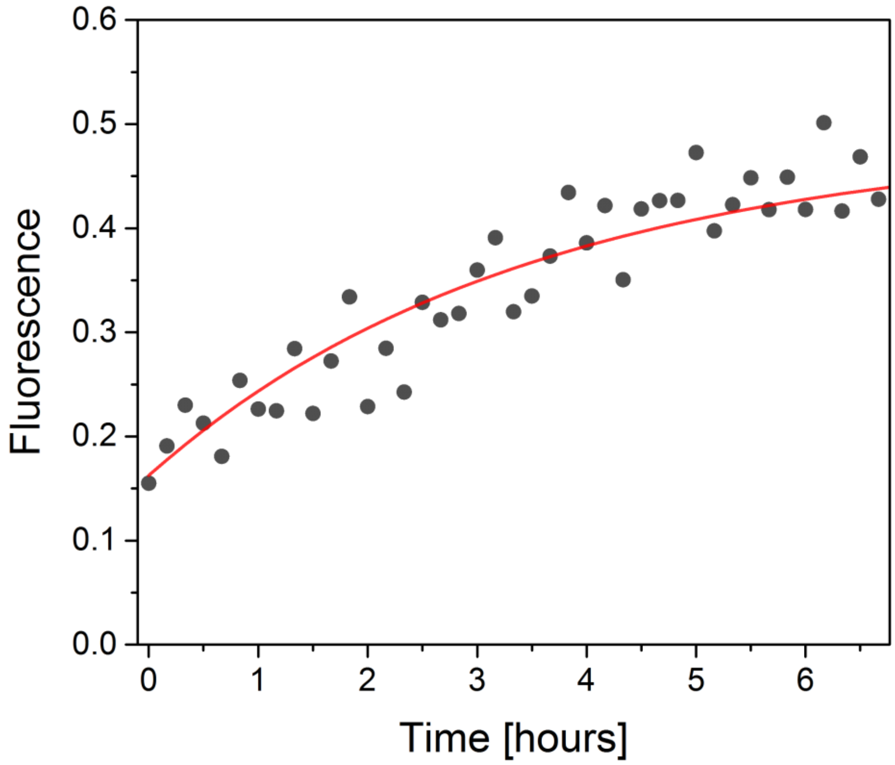


**Kinetic of red mEos4b thermal recovery after off-switching.**

Red mEos4b was illuminated for 170 seconds with 561-nm light (2 $kW/cm^2$), inducing photoswitching, and was subsequently left in the dark for 6 hours. The red fluorescence level was monitored by using short pulses (50 ms) of 561-nm light at low power density (0.2 $kW/cm^2$) every 10 minutes, so as to limit the influence of the readout light on the observed fluorescence recovery. A recovery half time of ~2 hours was fitted using a monoexponential model (red line).

**Supplementary Figure 5.**

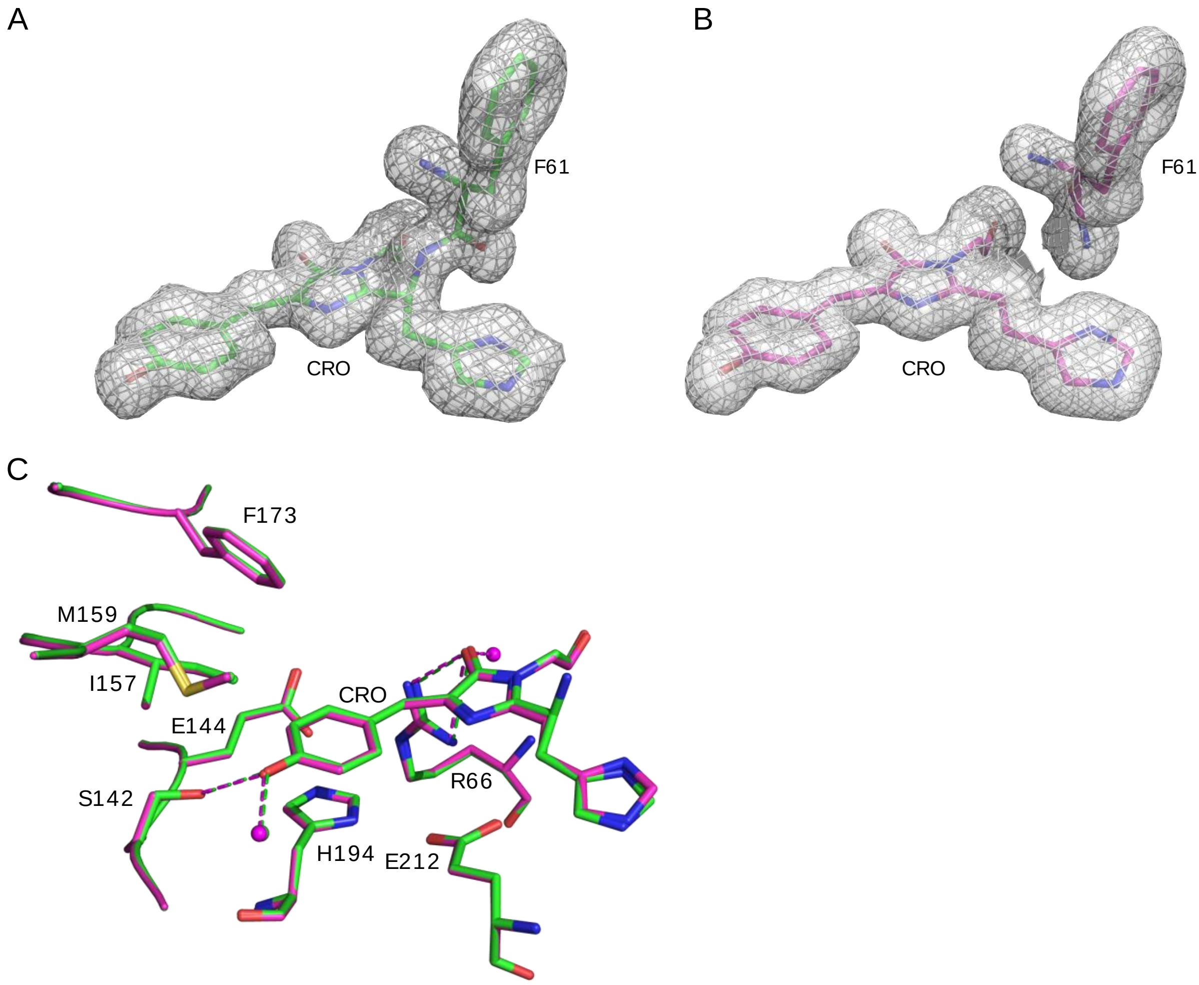


**Green and photoconverted red state crystal structure of mEos4b.**

(A) View of the chromophore and Phe61 in the green state before photoconversion in the $F_{obs}$-$F_{calc}$ omit electron density map (gray) contoured at 3 r.m.s.d. (B) Same view in the photoconverted red state. The cleavage between the chromophore and the preceding phenylalanine 61 is clearly revealed (C) Superposition of the green and photoconverted red mEos4b crystal structures, depicted in green and magenta sticks, respectively. Spheres represent water molecules; hydrogen bond are represented with dashes.

**Supplementary Figure 6.**

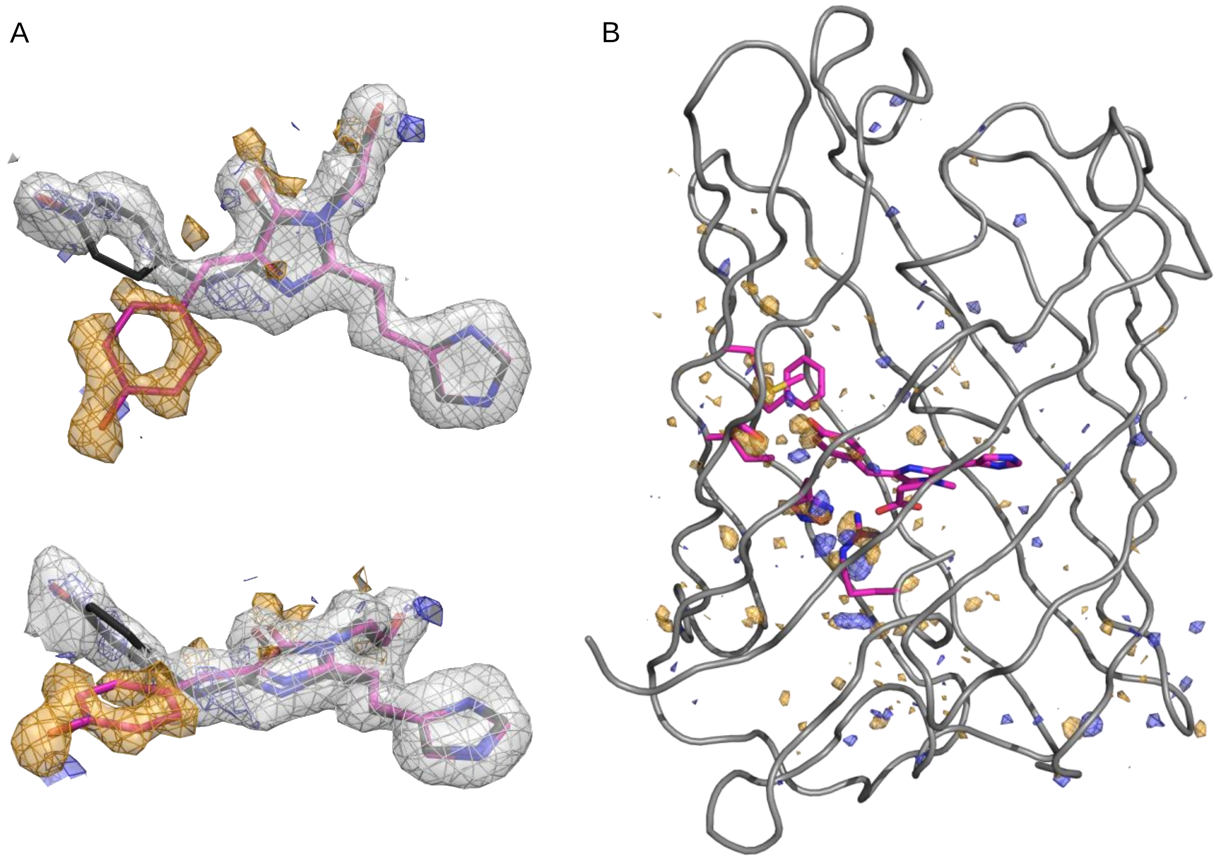


**Electron density maps of the bright and long-lived dark state of red mEos4b.**

(A) Top and front view of the bright (magenta) and longer-lived dark state $Red_{off}$ (black) chromophore of red mEos4b in the extrapolated $2F_{obs,extr.illuminated}$-$F_{calc,illuminated}$ (coloured in gray, contoured at 1 r.m.s.d.) and experimental difference $F_{obs,illuminated} - F_{obs,non\text{-}illuminated}$ electron density maps contoured at -3 r.m.s.d. (orange) and +3 r.m.s.d. (blue). (B) Experimental difference $F_{obs,illuminated} - F_{obs,non\text{-}illuminated}$ electron density map of the whole protein contoured at ±4 r.m.s.d. (orange and blue). See Online Methods for details on density maps calculations.

**Supplementary Figure 7.**

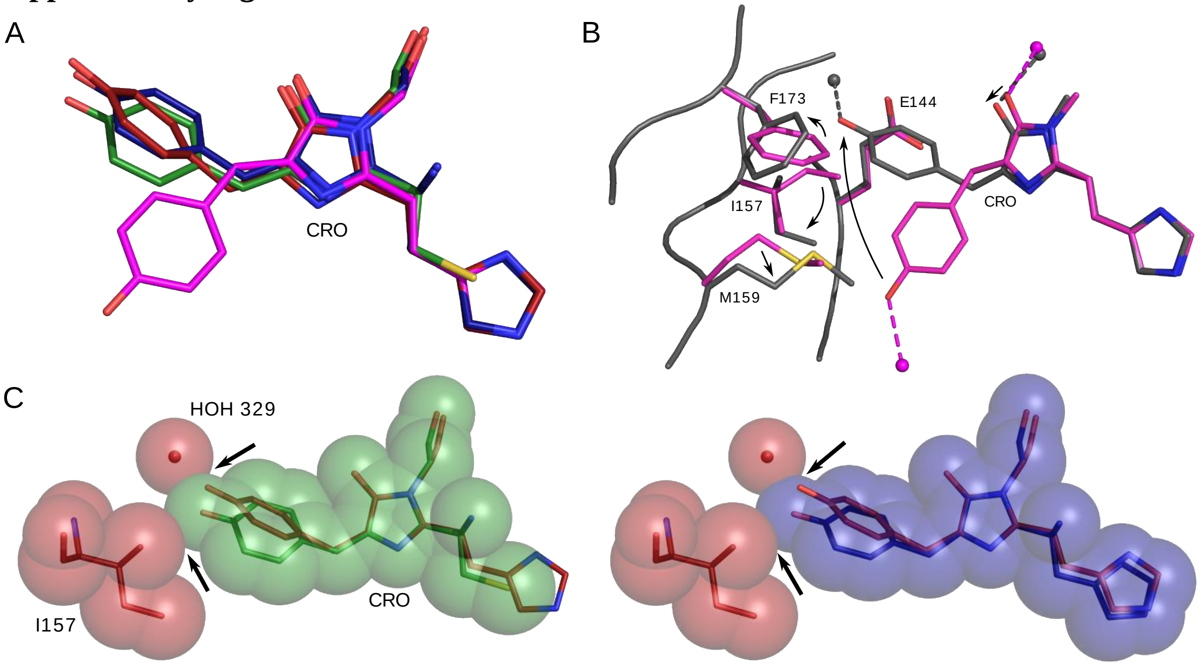


**The dark state chromophore adopts a frustrated *trans* conformation.**

(A) Chromophore superposition of red mEos4b (magenta), its long-lived dark state $Red_{off}$ (dark red), Dronpa in its switched-off state (green, PDB ID: 2POX) and IrisFP in its green switched-off state (blue, PDB ID: 2VVI). (B) Structural changes that lead to the frustrated *trans* configuration of the chromophore in red mEos4b. Arrows point to chromophore or neighboring residues motions (C) Views of hypothetical *trans* Dronpa (green) or green IrisFP (blue) chromophores with their imidazolinone moieties superposed onto that of red mEos4b in its long-lived dark state (dark red). The Dronpa and IrisFP chromophores are depicted in both ball-and-stick and van der Waals spheres representations, as well as Ile157 and water 329 from mEos4b. The van der Waals spheres suggest steric clashes of the "non-frustrated" Dronpa or IrisFP chromophores with Ile157 or water 329 (arrows).

**Supplementary Figure 8.**

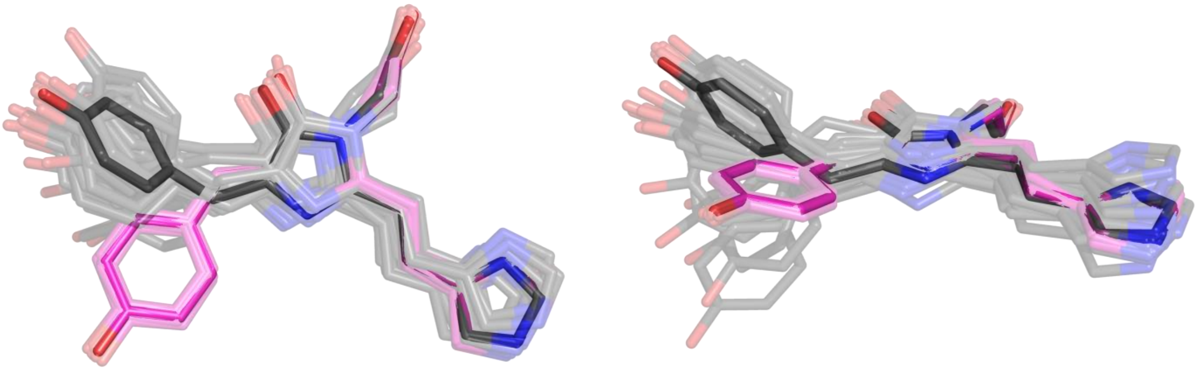

**Ensemble refinement of the bright and dark state mEos4b chromophore.**

Top and front view of the red mEos4b chromophore in its bright (magenta) and long-lived dark state (gray). The respectively 52 and 34 chromophore models derived from ensemble refinement are shown.

**Supplementary Figure 9.**

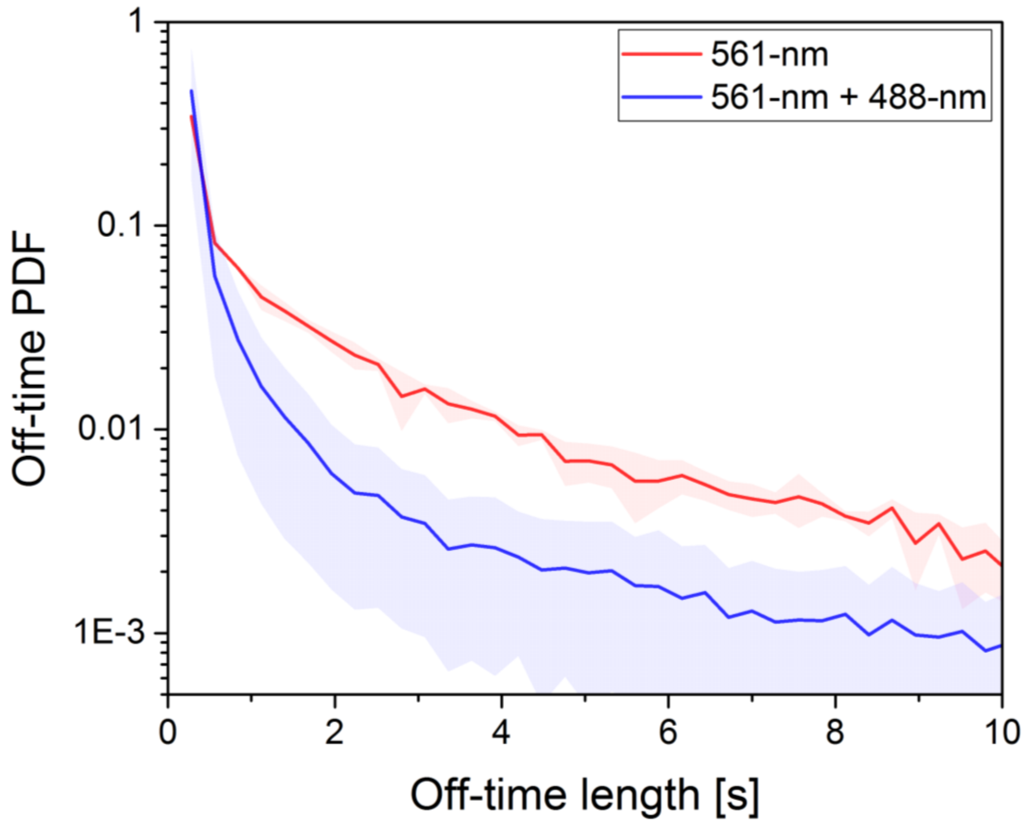


**Intermittency histogram of mEos4b fused to the CD86 receptor in a fixed HeLa cell.**

A CD86-mEos4b fusion protein was expressed in HeLa cells, and imaged under either 561-nm light (0.6 kW/cm$^2$) alone, or 561- + pulsed 488-nm light (10 W/cm$^2$, 10% duty cycle). As observed *in vitro*, illumination with 488-nm light drastically reduces the number of long intermittencies and increases the number of short intermittencies. PDF: probability density function.

**Supplementary Figure 10.**

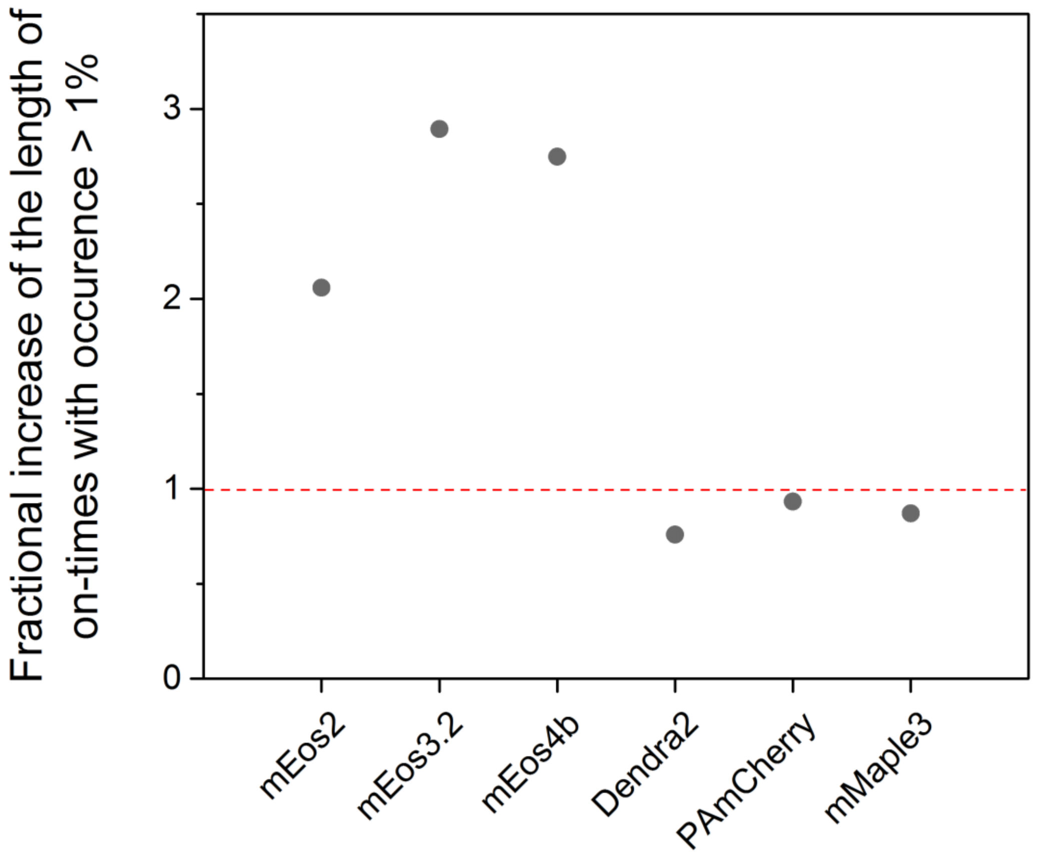


**Increase of on-times in the presence of 488-nm light for different popular FPs used in PALM.**

PALM datasets were acquired for 6 different photoactivatable or photoconvertible FPs under sole 561-nm light (0.5 kW/cm$^2$), or combined 561- and 488-nm light (120 W/cm$^2$, pulsed mode). The three tested Eos variants (mEos2, mEos3.2 and mEos4b) showed a 2- to 3-times increase in the on-time duration above which the 1% longest on-times are registered, when 488-nm illumination was used, whereas the other fluorescent proteins tested (Dendra2, PAmCherry and mMaple3) showed no improvement, possibly because less pronounced photoswitching or more pronounced photobleaching. The different behaviors of the mEos variants and Dendra2/mMaple3 might be connected to their different photophysical behavior as described in ref[1].

**Supplementary Figure 11.**

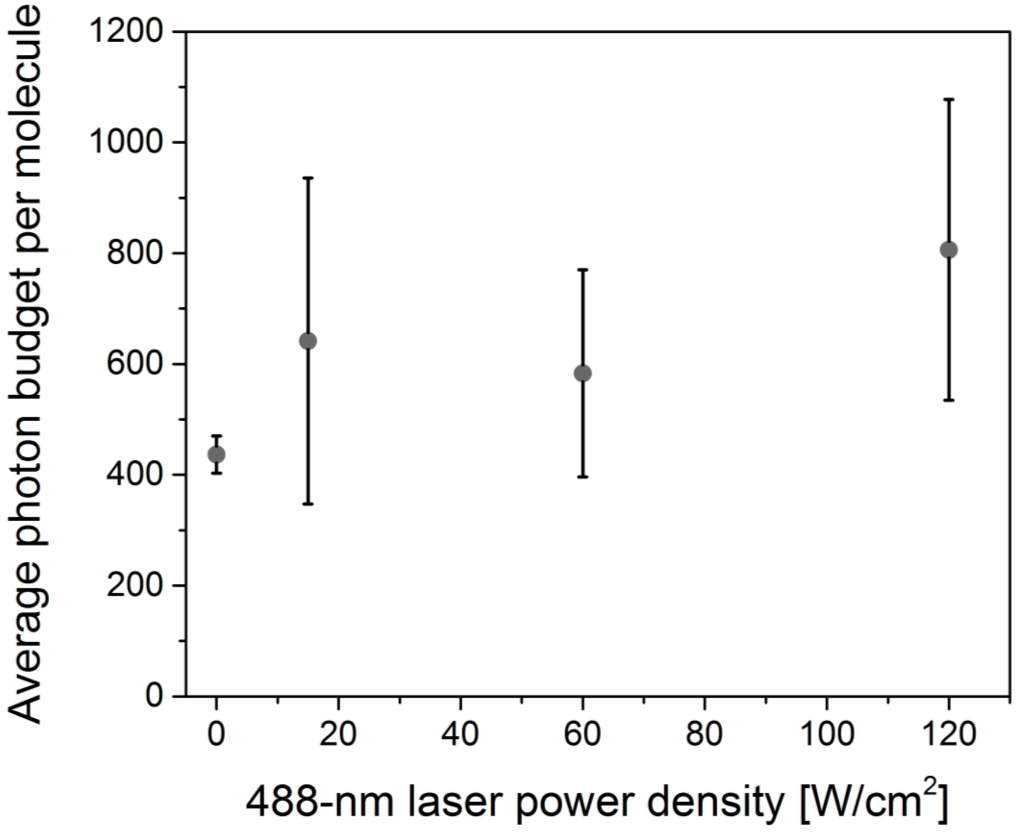


**Photon budget of single red mEos4b molecules under combined 561- and 488-nm illumination.**

Measured average photon budget under 0.5 kW/cm² of 561-nm light as a function of additional 488-nm illumination (10% duty cycle). Means and standard deviations from triplicate experiments are plotted. Increasing the 488-nm illumination power-density does not significantly affect the photon budget of red mEos4b molecules, showing that, at these intensity levels, 488-nm light does not increase substantially their photobleaching rate.

**Supplementary Figure 12.**

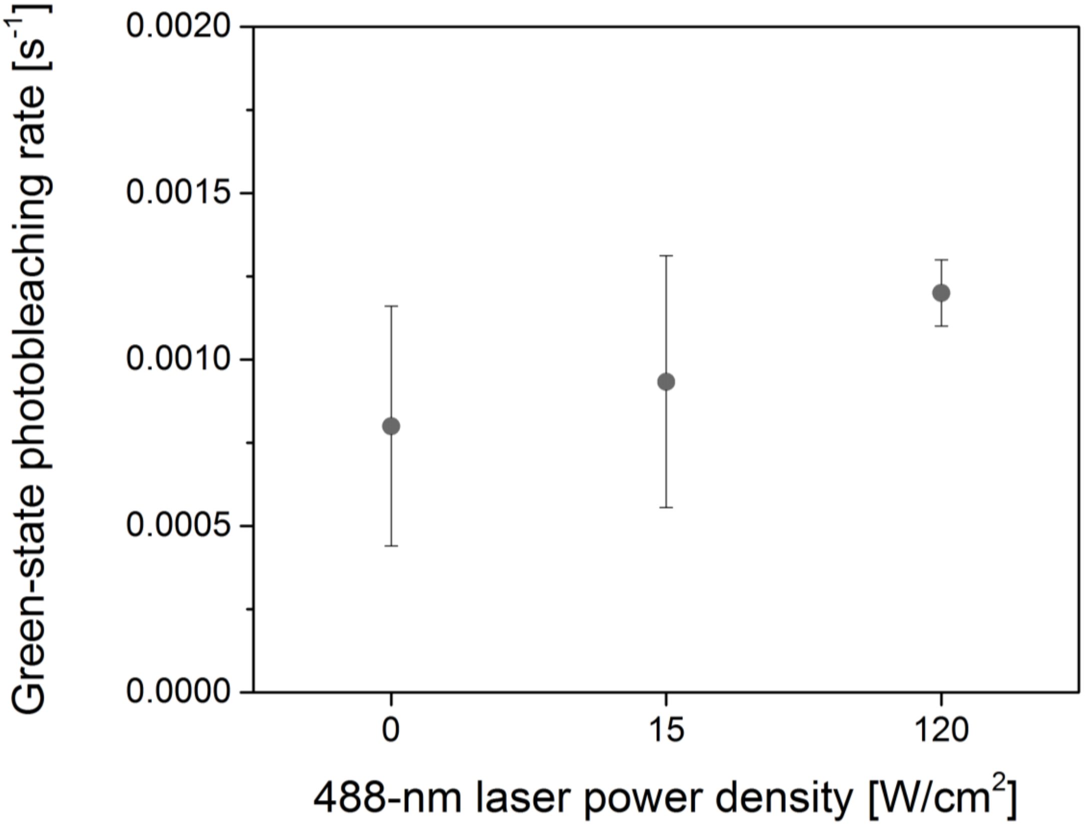


**Illumination at 488-nm during a PALM experiment induces minor photobleaching of green mEos4b.**

Ensemble fluorescence experiments were conducted on the green form of mEos4b using the same laser scheme and power densities as for PALM experiments (Fig. 2A), and with short pulses of weak 488-nm light (15 W/cm$^2$ for 10 ms every second) for fluorescence readout. Green-state photobleaching rates were retrieved as described in Supplementary Note 4. Upon addition of 488-nm light, only little additional green-state photobleaching was observed (photobleaching rate: 0.0012 s$^{-1}$ at the highest 488-nm laser power density, against 0.0008 s$^{-1}$ in the absence of 488-nm light).

**Supplementary Figure 13.**

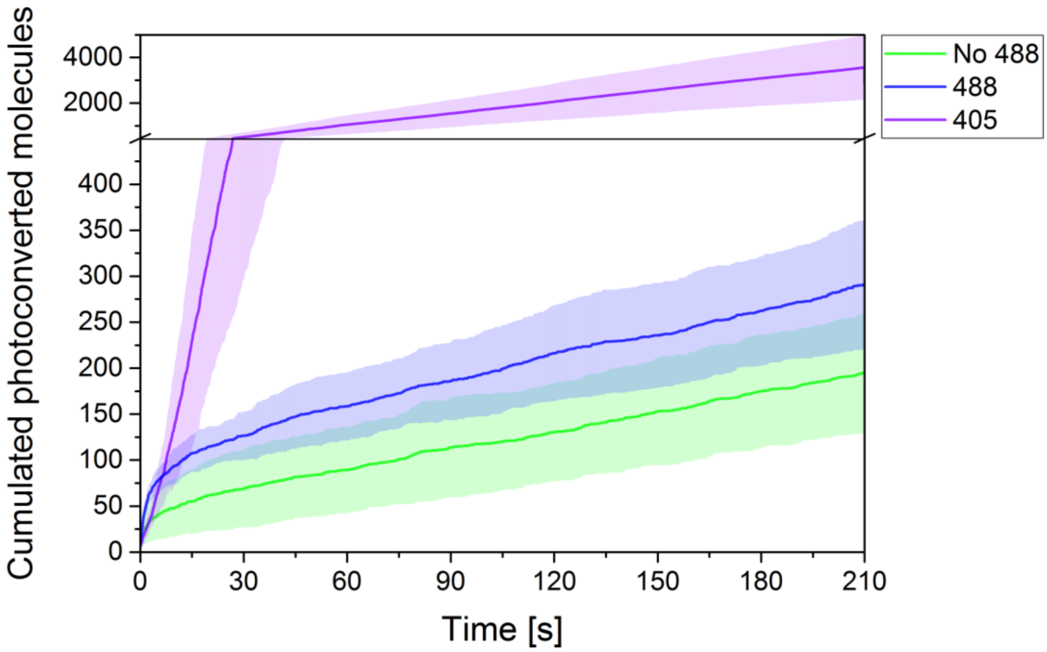


**Additional photoconversion due to 488-nm illumination.**

PALM experiments were conducted on PAA-embedded mEos4, under either sole 561-nm readout light (0.5 kW/cm$^2$), with combined 561- and 488-nm light (120 W/cm$^2$, 1/10$^{th}$ of the frametime), or with combined 561- and 405-nm light (0.6 W/cm$^2$, 1/10$^{th}$ of the frametime). 488-nm illumination induces slightly accelerated photoconversion, on a level however much lower than the 405-nm illumination typically used in PALM (note the Y-axis break at 500).

**Supplementary Figure 14.**

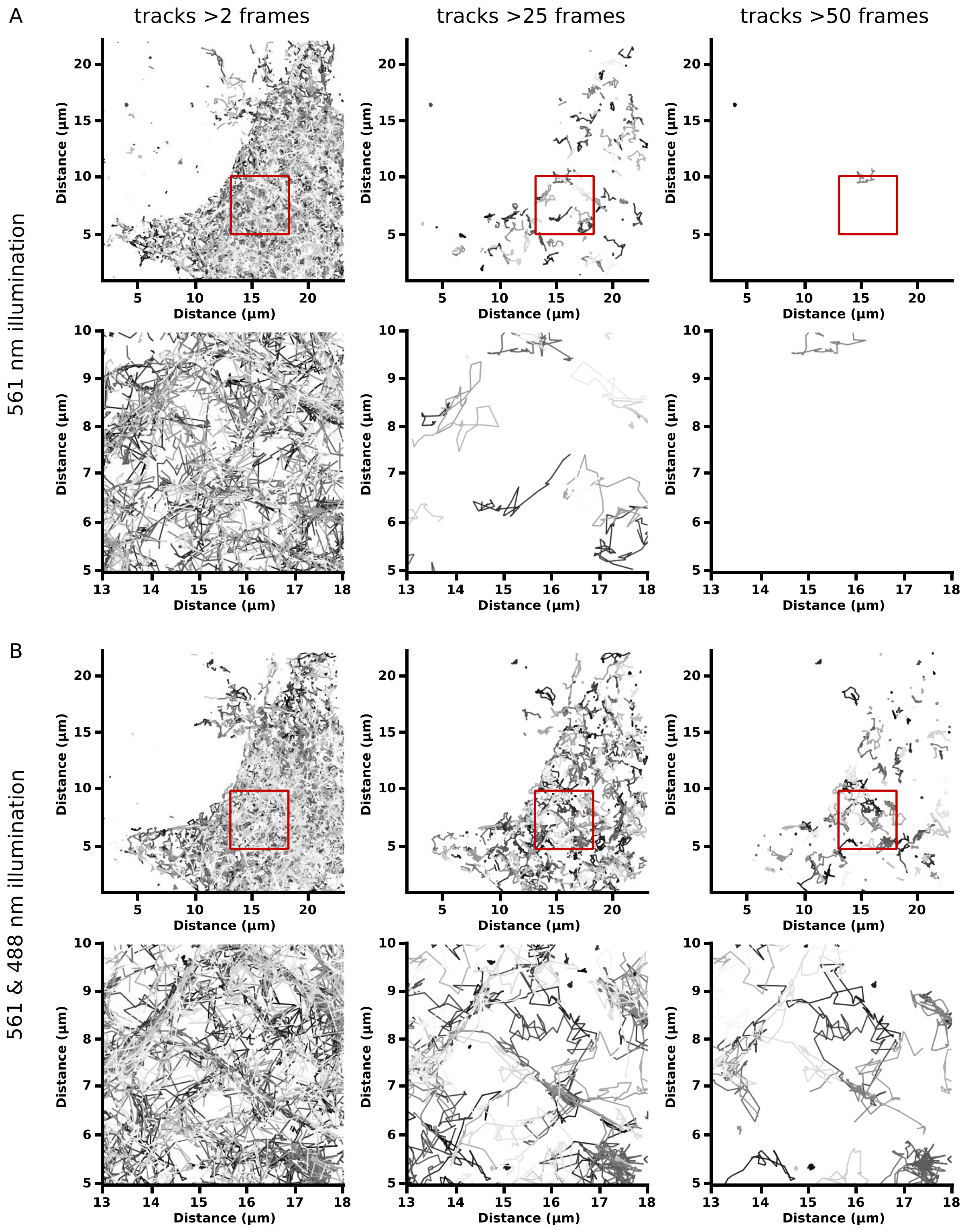


**Influence of 488nm illumination during single particle tracking.**

MAP4-mEos4b-labeled COS-7 live cell spt-PALM imaging without (A) and with (B) additional 488-nm illumination (4.8 W/cm2). The columns show respectively all tracks longer than 2 frames (0.08 s), 25 frames (1 s) and 50 frames (2 s). The red square indicates the region that has been zoomed in the bottom part.

**Supplementary Figure 15.**

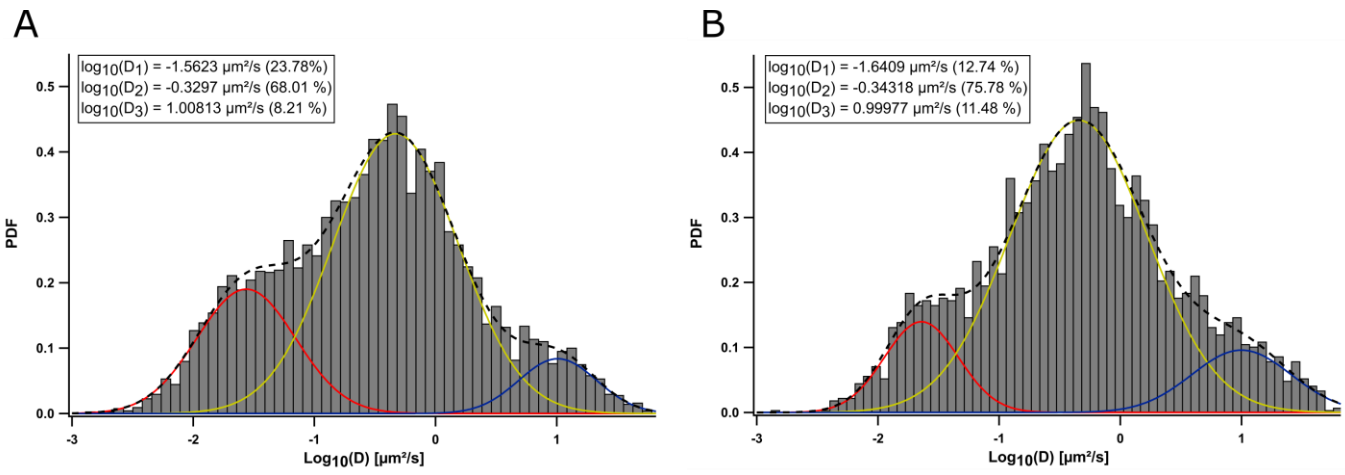


**Probability density function of the $log_{10}$-transformed diffusion coefficients from all MAP4-mEos4b tracking experiments with a length > 0.8 s, in the absence (A) or presence (B) of additional 488-nm illumination**

The two curves of the distribution of the $log_{10}$-transformed diffusion coefficients, extracted from the HMM-Bayes software package, show the clear presence of three different diffusion regimes. Within the graph, the three different distributions are shown, each with their respective contributions. The differences between the two distributions are statistically nonsignificant (Residual differences (see inset) are statistically nonsignificant ($\chi^2$ test with $\alpha$ = 0.01).

**Supplementary Figure 16.**

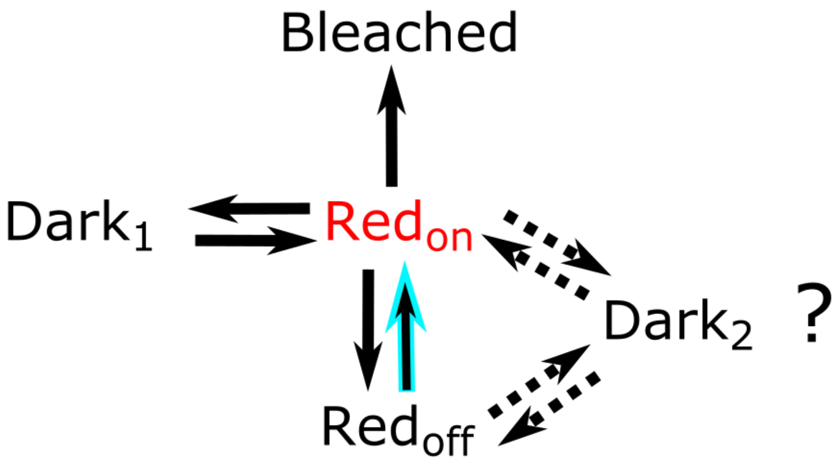


**Tentative photophysical scheme for red mEos4b.**

After photoconversion (not represented) the red-fluorescent form of mEos4b ($Red_{on}$) can either photobleach, enter a light-insensitive short-lived dark-state ($Dark_1$), or enter a light-sensitive longer-lived dark (switched-off) state ($Red_{off}$). From there, or from $Red_{on}$, it might be able to enter a further uncharacterized dark-state ($Dark_2$), relaxing towards either $Red_{on}$ or $Red_{off}$. Illumination at 488-nm allows a faster recovery from the $Red_{off}$ to the $Red_{on}$ state (cyan arrow).

**Supplementary Figure 17.**

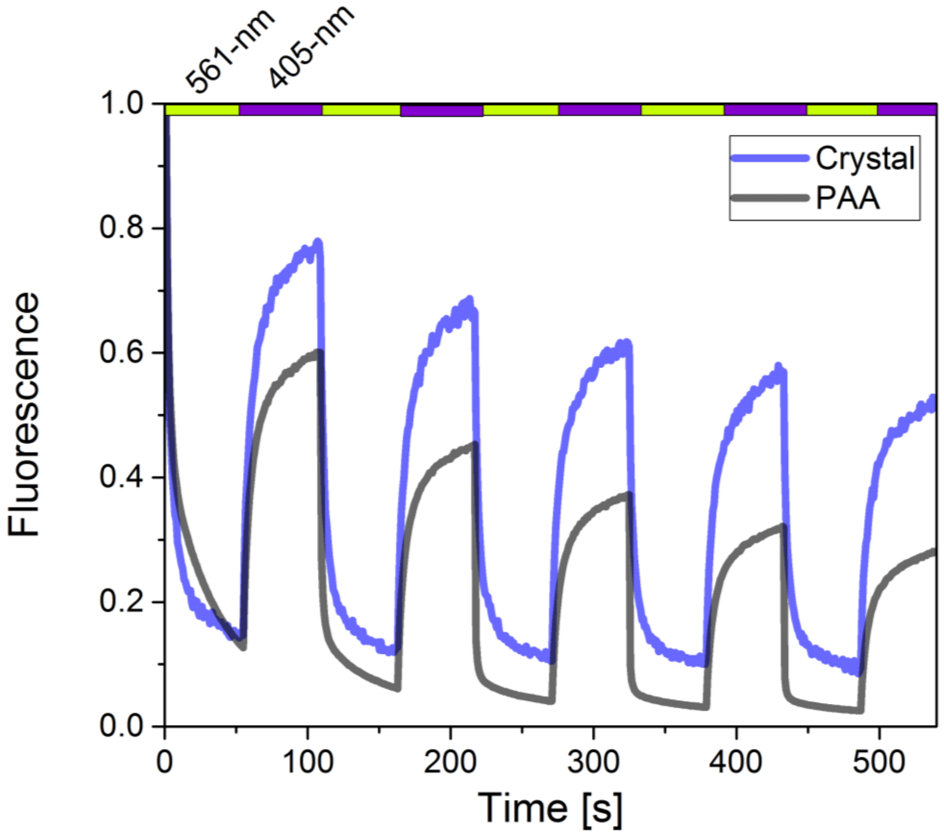


**PAA embedded and *in crystallo* red mEos4b reversible photoswitching.**

Reversible photoswitching of mEos4b embedded in PAA (black curve) or *in crystallo* (blue curve) under alternating 561- and 405-nm light (green and violet bar shown on top, respectively 70 and 0.3 W/cm2 for *in crystallo* data, and 25 and 1 W/cm2 for PAA data).

**Supplementary Figure 18.**

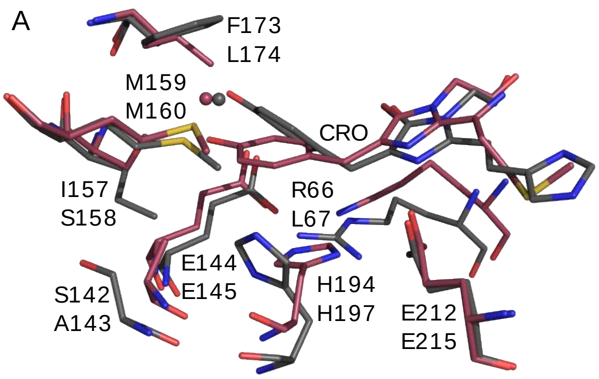


**Superposition of the dark chromophore and its environment in red mEos4b and reversibly switchable red fluorescent proteins.**

Superposition of red mEos4b in the long-lived dark state (gray) with (A) asFP595 (PDB ID 2A50, dark red) and (B) rsTagRFP (PDB ID 3U8A, dark red) in their respective switched-off states. In all cases, the chromophore of mEos4b is found significantly more distorted.

**Supplementary Figure 19.**

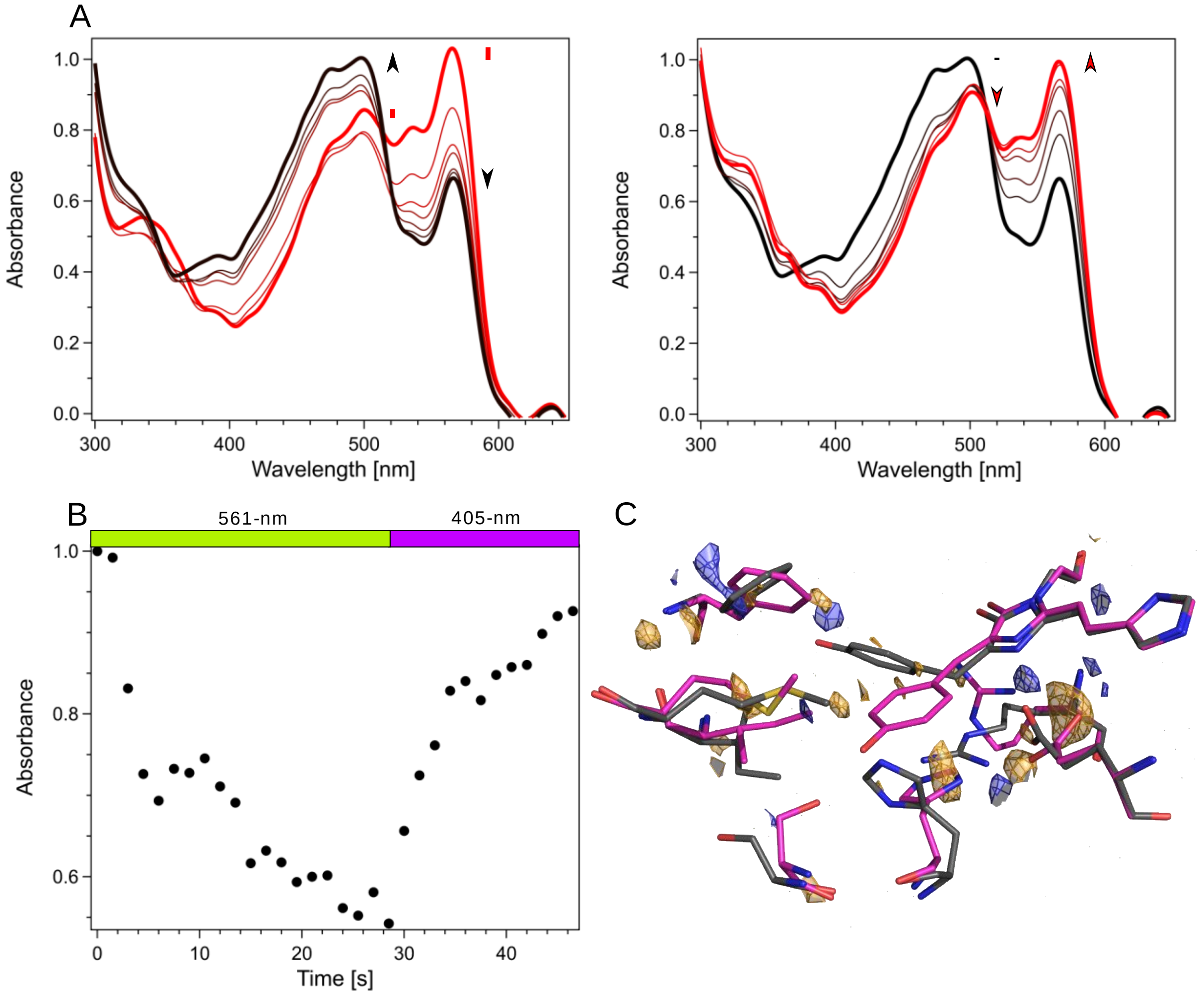


**Reversibility of the dark state trapped in crystalline mEos4b.**

(A) Evolution of normalized absorption spectra of a control mEos4b crystal upon sequential illumination with 561-nm laser light (left) and 405-nm light (right). Arrows show the evolution of absorption bands. Recovery of the red fluorescent state is evident upon 405 nm light illumination (right). (B) Time profile of the absorption band of the anionic chromophore during 561-nm illumination (green bar) and subsequent 405-nm illumination (violet bar). (C) Difference electron density map $F_{obs,Side1}$-$F_{obs,Side2}$ between the treated (side 1) and untreated (side 2) parts of the crystal, contoured at -3 r.m.s.d. (orange) and +3 r.m.s.d. (blue) on top of the red bright and long-lived dark state crystal structures. The absence of significant difference electron density on the chromophore suggests that mEos4b molecules have mostly returned to the red fluorescent state.

**Supplementary Figure 20.**

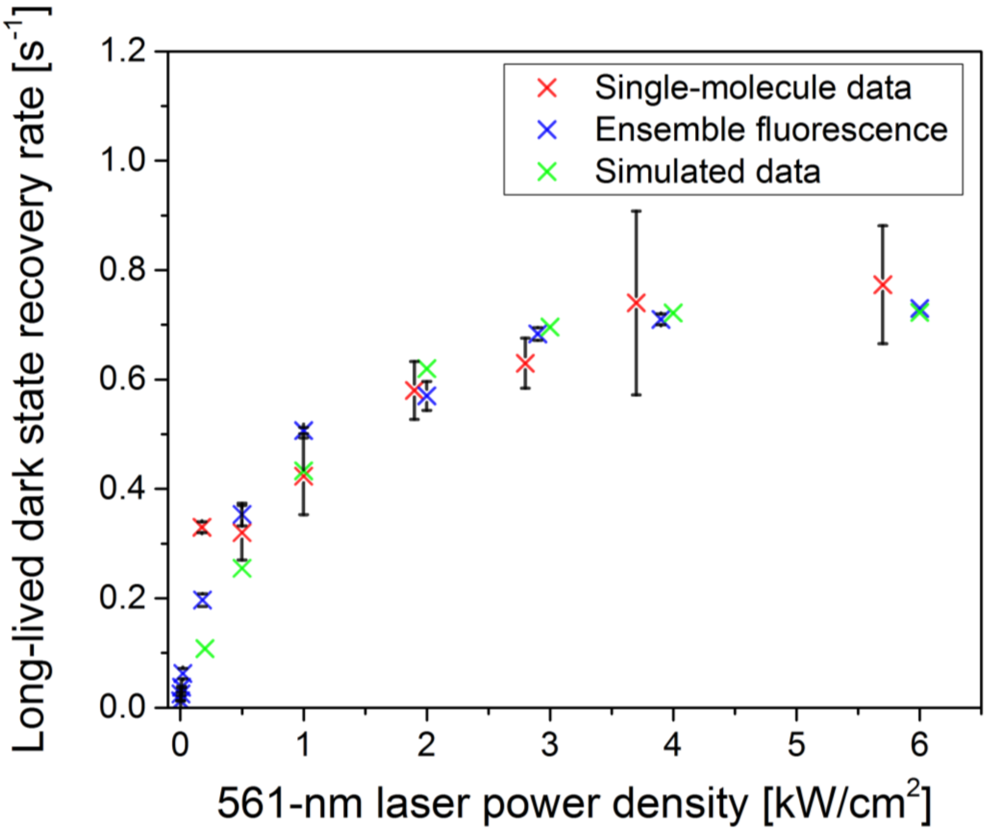


**Simulation of the rate saturation effect upon 561-nm laser power titration.**

Ensemble fluorescence data were simulated based on the kinetic model of Supplementary Fig. 15, and fitted with the same simplified kinetic model (i.e. assuming only two dark states) as used for the analysis of ensemble experiments. By adjusting the rate constants of the model (Supplementary Table 4), a recovery rate saturation behavior very similar to the one observed experimentally could be reproduced.

**Supplementary Table 1.** Data collection and refinement statistics of the bright and dark-state crystal structures of mEos4b.

| | **Green mEos4b** | **Red mEos4b bright state (*Red<sub>on</sub>*)** | **Red mEos4b long-lived dark state (*Red<sub>off</sub>*)** |
|---|---|---|---|
| **PDB ID** | 6GOY | 6GP0 | 6GP1 |
| Space group | $P2_12_12_1$ | $P2_12_12_1$ | $P2_12_12_1$ |
| Unit cell parameters | | | |
| a (Å) | 38.60 | 38.61 | 38.60 |
| b (Å) | 58.09 | 57.90 | 57.96 |
| c (Å) | 103.27 | 102.64 | 102.49 |
| Resolution range (Å) | 51.64 – 1.65 (1.69 – 1.65)* | 36.14 – 1.50 (1.54 – 1.50) | 36.13 – 1.30 (1.33 – 1.30) |
| No. of unique reflections | 27933 (2083) | 36532 (2643) | 56087 (4069) |
| $R_{meas}$ (%) | 12.1 (62.5) | 5.9 (69.1) | 5.4 (57.7) |
| $\langle I/\sigma(I)\rangle$ | 6.59 (1.82) | 13.87 (1.79) | 16.52 (2.14) |
| Completeness (%) | 97.2 (99.0) | 96.3 (95.4) | 97.7 (96.8) |
| Multiplicity (%) | 2.75 (2.74) | 2.45 (2.40) | 3.60 (2.75) |
| No. of molecules per AU | 1 | 1 | 1 |
| $R_{work}/R_{free}$† (%) | 16.78 / 20.59 | 15.55 / 19.00 | 24.89 / 30.43# |
| Rmsd | | | |
| Bond length (Å) | 0.005 | 0.004 | 0.007 |
| Bond angles (°) | 0.801 | 0.866 | 0.941 |
| No. of protein atoms | 1903 | 1957 | 1875 |
| No. of waters | 269 | 347 | 345 |
| Average B-factor (Å2) | | | |
| Main/side chain | 15.68 / 19.42 | 15.0 / 18.9 | 13.8 / 7.0 |
| Waters/Ligands | 30.66 / 38.27 | 31.7 / NA | 26.6 / NA |
| Ramachandran statistics (%) | | | |
| Favored | 99.1 | 99.5 | 99.1 |
| Outliers | 0.0 | 0.0 | 0.00 |

* Values in the parentheses refer to the highest resolution shell

† $R_{free}$ is calculated using a 0.05 % fraction of random reflections excluded from refinement

\# Structure refinement using extrapolated structure factors

**SupplementaryTable 2.** Data collection and refinement statistics of control crystal structures of mEos4b. On side of the crystal was treated with subsequent 561- and 405 nm illumination, whereas the other side was kept non-illuminated.

| | **Red mEos4b Side 1** | **Red mEos4b Side 2** |
|---|---|---|
| Space group | $P2_12_12_1$ | $P2_12_12_1$ |
| Unit cell parameters | | |
| a (Å) | 38.57 | 38.48 |
| b (Å) | 57.74 | 57.64 |
| c (Å) | 102.56 | 102.52 |
| Resolution range (Å) | 51.29 – 1.65 (1.69 – 1.65) | 51.27 – 1.70 (1.74 – 1.70) |
| No. of unique reflections | 28181 (1934) | 25746 (1831) |
| $R_{meas}$ (%) | 7.8 (63.8) | 9.0 (58.0) |
| $\langle I/\sigma(I) \rangle$ | 17.05 (3.04) | 16.39 (3.24) |
| Completeness (%) | 99.3 (92.9) | 99.5 (97.1) |
| Multiplicity (%) | 5.36 (5.24) | 5.37 (5.29) |
| No. of molecules per AU | 1 | 1 |

* Values in the parentheses refer to the highest resolution shell

**Supplementary Table 3.** Average chromophore torsion and bond angles with standard deviation when more than one monomer is present in the asymmetric unit.

| **Protein and state** | **PDB ID** | **Tilt# (º)** | **Twist# (º)** | **Methylene bridge (º)** |
|---|---|---|---|---|
| mEos4b red bright& | 6GP0 | -165.2 | -21.0 | 127.0 |
| mEos4b red long-lived dark& | 6GP1 | -28.5 | 83.2 | 123.4 |
| IrisFP green-on | 2VVH | -175.4 ± 1.7 | -12.0 ± 0.6 | 128.7 ± 0.9 |
| IrisFP green-off | 2VVI | -3.6 ± 0.7 | 40.3 ± 6.4 | 125.4 ± 0.5 |
| IrisFP red-on state | 2VVJ | -165.3 ± 2.1 | -21.6 ± 2.4 | 127.6 ± 0.6 |
| IrisFP red-off | * | 2.8 ± 3.9 | 41.3 ± 3.1 | 125.4 ± 1.4 |
| Dronpa green-on | 2IE2 | -178.7 ± 0.3 | -7.0 ± 1.2 | 129.5 ± 0.3 |
| Dronpa green-off | 2POX | -20.7 ± 4.9 | 40.1 ± 6.3 | 133.3 ± 1.0 |
| pcDronpa green-on | 4HQ8 | -168.5 ± 1.0 | -17.0 ± 1.9 | 130.5 ± 1.3 |
| pcDronpa green-off | 4HQ9 | -24.3 ± 7.9 | 57.3 ± 5.4 | 127.8 ± 0.8 |
| rsTagRFP red-on | 3U8C | -179.1 ± 1.1 | -5.6 ± 1.6 | 132.5 ± 4.1 |
| rsTagRFP red-off | 3U8A | -16.7 ± 2.1 | 6.7 ± 1.9 | 137.8 ± 0.7 |
| mTFP0.7 cyan-on | 2OTB | -165.2 ± 0.8 | -16.9 ± 2.7 | 127.2 ± 0.8 |
| mTFP0.7 cyan-off | 2OTE | -16.7 ± 1.3 | 66.8 ± 0.3 | 132.1 ± 1.6 |
| asFP595 red-on | 2A52 | -162.8 ± 2.5 | -15.2 ± 3.7 | 127.9 ± 1.9 |
| asFP595 red-off | 2A50 | -14.2 ± 0.4 | 29.5 ± 0.3 | 128.8 ± 1.6 |

& Only one monomer in asymmetric unit

* The Iris red-off structure was not experimentally determined but proposed using computational means.

### Tilt and Twist torsion angles as described in Quillin *et al.*[2] with the tilt angle calculated over C12-C9-C8-C7 atoms and twist angle over C9-C8-C7-C6, as shown below.

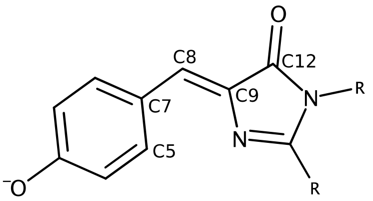


**Supplementary Table 4.** Rate constants used for simulation of ensemble level experiments based on the photophysical scheme of Supplementary Fig. 15 (for a 561-nm laser power density of 1 kW/cm$^2$).

| **Process** | **Rate [s$^{-1}$]** | **Relaxation** |
|---|---|---|
| Red$_{on}$ bleaching | 0. 1 | Light-induced |
| Red$_{on}$ to Dark$_{on}$ | 0.5 | Light-induced |
| Dark$_{on}$ to Red$_{on}$ | 16 | Thermal |
| Red$_{on}$ to Red$_{off}$ | 0.5 | Light-induced |
| Red$_{off}$ to Red$_{on}$ | 0.3 | Light-induced |
| Red$_{off}$ to Dark$_{off}$ | 0.3 | Light-induced |
| Dark$_{off}$ to Red$_{on}$ | 0.6 | Thermal |

#### Supplementary Discussion

**Note 1. Fitting of fluorescence intermittencies histograms.**

Upon fitting of the mEos4b intermittency histograms using a bi-exponential model accounting for the two previously characterized dark-states of red mEos variants[3,4] (Supplementary Fig. 1A), we observed that the fit did not account well for the observed long off-times (> 1 second). This deviation of the model might be partially explained by spurious effects such as limited accuracy of single-molecules detection and localization, or laser beam heterogeneities. Alternatively, or in addition, as detailed below in Supplementary Note 6, it could also result from a more complex photophysical scheme involving more than two dark-states (Supplementary Fig. 15). A tri-exponential model yields a very satisfactory fit of the intermittencies histograms (Supplementary Fig. 1B). However, whatever the cause for the deviation from a bi-phasic behavior, the recovery rate of $Red_{off}$ cannot be directly deduced from the two slowest rates fitted with this tri-exponential model.

**Note 2. Ensemble-level dark-state accumulation in solution and *in-crystallo.***

By conducting illumination experiments at the ensemble level on mEos4b embedded in a polyacrylamide gel or in the crystalline form (Fig. 1C, Supplementary Fig. 3, Supplementary Fig. 17), a very similar behavior is observed, confirming that mEos4b is still able to enter the longer-lived dark state *in-crystallo*.

The exact kinetics are however slightly modified, which can be attributed to different factors. First, the density of proteins in a crystal is much higher than in a gel, giving rise to inner filtering effects. The high optical density creates heterogeneities, since proteins at the top of the crystal receive more photons than those at the bottom, which are shielded by the top layers. Another effect of crystal packing is that the chromophore dipoles are not randomly oriented, but have four main orientations within space group $P2_12_12_1$. This also creates heterogeneities, with populations of favorably oriented chromophores efficiently absorbing light, and populations of unfavorably oriented chromophores showing reduced absorption.

**Note 3. Structural basis for the frustrated *trans* conformation of the mEos4b chromophore in the long-lived dark state.**

The crystal structure of the red non-illuminated part of the crystal contained a mixture of both the green and red state, as could be deduced by the presence of weak electron density between the chromophore and Phe61. Therefore, the green state crystal structure was determined first.

The crystal structure of red mEos4b shows the typical pattern of other photoconverted EosFP derivatives described earlier[5] with the backbone break of the $C_\alpha$-N bond of His62 associated to an extended $\pi$-conjugation of the chromophore being the only main difference as compared to the green mEos4b structure (Supplementary Fig. 5).

The crystallographic data suggest that the longer-lived dark state in red mEos4b ($Red_{off}$) is essentially the consequence of *cis-trans* isomerization. Indeed, the q-weighted $F_{obs,illuminated}$– $F_{obs,non\text{-}illuminated}$ difference electron density map of Fig. 1D and Supplementary Fig. 6 indicate a strong loss of density at the *cis* isomeric position of the chromophore, a gain of density at the *trans* position, and substantial conformational changes of neighboring amino acids, notably Arg66, Ser142, Ile157, Phe173, His194 and Glu212, that closely resemble those occurring in anthozoan-derived green RSFPs like Dronpa[6], pcDronpa[7], IrisFP[8] and mTFP0.7[9] (Fig. 1E). The rather weak and diffuse gain in electron density in the *trans* state however indicates that the chromophore has increased flexibility compared to the bright state (Supplementary Fig. 6 and 8). This increase in flexibility is much more pronounced than in all structures of switched off green RSFPs reported thus far. Furthermore, the chromophore displays significantly bigger tilt and twist dihedral angles as well as a decreased methylene bridge bond angle (Supporting Table 3), resulting in a strong overall distortion while likely maintaining electron conjugation between the hydroxybenzylidene and imidazolinone cyclic moieties. As a consequence of the chromophore distortion, the stabilizing interaction with Glu144 typically seen in switched-off green RSFPs is weakened, but is complemented by a hydrogen bond with a water molecule (Supplementary Fig. 7C). We propose that the sub optimal set of interactions maintained by the mEos4b chromophore with its environment in the long-lived dark state forms the basis for a poor switching contrast, i.e. the capacity of 561-nm photons to promote recovery to the red fluorescent state relatively efficiently, providing the blinking behavior observed in PALM. Of note, in red RSFPs for which crystallographic structures of the switched state have been reported, such as asFP595[10] or rsTagRFP[11], the strong chromophore distortion found in mEos4b is not observed (Supplementary Fig. 18).

A remarkable difference between mEos4b and other RSFPs in their isomerized dark states is a small shift in the chromophore's position towards strands 7 and 8 of the $\beta$-barrel, clearly visible at the level of the imidazolinone ring (Supplementary Fig. 7A). We hypothesize that this shift may form the basis for the observed chromophore bending, as a relatively flat *trans* chromophore as observed in Dronpa or IrisFP would result in a steric conflict, notably with Ile157 (Supplementary Fig. 7C). To accommodate this proximity with Ile157, the chromophore bends and causes a cascade of small rearrangements in which Phe173 and Met159 need to deviate from their preferred side chain conformation (Supplementary Fig. 7B).

Based on these structural observations, we performed site-saturation mutagenesis at positions 157 and 173, in the hope that some combinations of residues at these positions could prevent the chromophore to find a stable conformation in the *trans* state. However, amongst those mutants that correctly maturated and could be successfully photoconverted, we could not find one exhibiting significantly reduced slow-blinking as compared to mEos4b itself (data not shown).

One may wonder whether the crystallographic structure of the dark state described above could be contaminated by irreversible photobleaching. As a control experiment, we illuminated one side of a crystal sequentially with 561-nm light followed by 405-nm recovery and calculated the q-weighted experimental difference map between both sides of the crystal (Supplementary Fig. 19). The absence of high electron density peaks indicates that the bright state structure can be recovered and that the crystal structure presented in Fig. 1D represents that of the longer-lived dark state.

**Note 4. Side effects of 488-nm illumination.**

A side-effect to the use of 488-nm light during PALM experiments is a modest increase of the photoconversion rate of the mEos4b molecules. The extent of this effect is shown in Supplementary Fig. 13. PALM datasets acquired in the presence of 488-nm light (120 W/cm$^2$, during 1/10$^{th}$ of the frametime) showed approximately 1.5-times more photoconverted molecules than datasets acquired under sole 561-nm light. However, this effect was still small compared to the more than 10-times increase in the number of photoconverted molecules obtained using 405-nm light (0.6 W/cm$^2$, 1/10$^{th}$ of the frametime).

The use of additional 488-nm light during PALM experiments raises the question of the photobleaching of mEos4b molecules. Increased photobleaching of the red molecules would lead to decreased track lengths and localization precision, while photobleaching of the green state might lead to incomplete photoconversion.

To assess red state photobleaching, the photon budget of single red mEos4b molecules embedded in PAA was measured under increasing 488-nm laser power density (Supplementary Fig. 11). It was found that, at the used power densities (15 to 120 W/cm$^2$), the photon budget is not affected by the addition of 488-nm light. Together with the lengthening of track lengths observed in sptPALM when using 488-nm light (Fig. 2E), this indicates that photobleaching of red mEos4b by 488-nm light is very marginal.

Evaluating green state photobleaching at the single-molecule level has proven challenging, since it would require photoconverting all mEos4b molecules present in a field of view, and assessing the proportion that could not reach the red state. Hence, green state photobleaching was investigated by conducting ensemble experiments on green mEos4b, using a protocol based on ref[12]. The protein was illuminated for 100 seconds using the same laser scheme as in PALM experiments (Fig. 2A, pulsed mode). Every second, a short pulse of 488-nm light (10 ms, 15 W/cm$^2$) was applied to read fluorescence. The fluorescence decay was fitted using a kinetic model including two reversible dark-states and photobleaching, and the photobleaching rate retrieved (Supplementary Fig. 12). Already in the absence of 488-nm illumination in the PALM laser scheme, some photobleaching occurs ($8 \times 10^{-4} \pm 4 \times 10^{-4}$ s$^{-1}$), corresponding to the previously characterized green-state photobleaching by 561-nm light[12]. Addition of weak (15 W/cm$^2$) or mild (120 W/cm$^2$) 488-nm light in the PALM laser scheme results in a small increase of the photobleaching rate, to $9 \times 10^{-4} \pm 4 \times 10^{-4}$ s$^{-1}$ and $1.2 \times 10^{-3} \pm 1 \times 10^{-4}$ s$^{-1}$, respectively. Green-state photobleaching due to the use of additional 488-nm light during PALM experiments is therefore minimal compared to the bleaching produced by the 561-nm readout light.

Furthermore, the photoconversion induced by 488-nm light allows decreasing the intensity of the 405-nm illumination, and hence decreasing photobleaching due to this laser, both on the green and red states of the protein. Phototoxic effects due to 405 nm light may also be alleviated.

**Note 5. Photophysical scheme of red mEos4b, and hypothesis of an additional dark state *Dark$_2$*.**

Several pieces of evidence suggest that red mEos4b (and probably the other mEos variants) may access more than two dark states: i) the fact that 488-nm light appears unable to suppress entirely all long-lived intermittencies in sptPALM experiments ; ii) the tri-phasic

shape of the intermittencies histograms (Supplementary Fig. 1) ; iii) the observed saturation of the recovery rate of $Red_{off}$ when only 2 dark states are assumed.
To substantiate the hypothetical photophysical scheme of Supplementary Fig. 16, we simulated ensemble experiments similar to those from which experimental data were reported in Fig. 1B and analyzed them in the same manner. By adjusting the rates entered in the model of Supplementary Fig. 16, we obtained a saturation effect very similar to the one observed experimentally (Supplementary Fig. 20). The rates obtained in this way were in full agreement with expected values (Supplementary Table 4). According to this scenario, the observed saturation can be mostly attributed to the presence of the additional $Dark_2$ state, which returns to the $Red_{on}$ state by a thermal process, thus producing an increasing shelving effect as the 561-nm laser is increased.
